# Automated EEG mega-analysis I: Spectral and amplitude characteristics across studies

**DOI:** 10.1101/409631

**Authors:** Nima Bigdely-Shamlo, Jonathan Touryan, Alejandro Ojeda, Christian Kothe, Tim Mullen, Kay Robbins

## Abstract

Significant achievements have been made in the fMRI field by pooling statistical results from multiple studies (meta-analysis). More recently, fMRI standardization efforts have focused on enabling the joint analysis of raw fMRI data across studies (mega-analysis), with the hope of achieving more detailed insights. However, it has not been clear if such analyses in the EEG field are possible or equally fruitful. Here we present the results of a large-scale EEG mega-analysis using 18 studies from six sites representing several different experimental paradigms. We demonstrate that when meta-data are consistent across studies, both channel-level and source-level EEG mega-analysis are possible and can provide insights unavailable in single studies. The analysis uses a fully-automated processing pipeline to reduce line noise, interpolate noisy channels, perform robust referencing, remove eye-activity, and further identify outlier signals. We define several robust measures based on channel amplitude and dispersion to assess the comparability of data across studies and observe the effect of various processing steps on these measures. Using ICA-based dipolar sources, we also observe consistent differences in overall frequency baseline amplitudes across brain areas. For example, we observe higher alpha in posterior *vs* anterior regions and higher beta in temporal regions. We also detect consistent differences in the slope of the aperiodic portion of the EEG spectrum across brain areas. In a companion paper, we apply mega-analysis to assess commonalities in event-related EEG features across studies. The continuous raw and preprocessed data used in this analysis are available through the DataCatalog at https://cancta.net.

## 1 Introduction

Data-pooling of fMRI (functional magnetic resonance imaging) studies has proven extremely valuable for increasing statistical power, assessing inter-subject variability, and determining the generalizability and reproducibility of predictions (Costafreda, 2009). fMRI *meta-analyses* typically combine studies based on coordinates of peak activations (coordinate-based meta-analysis, CBMA) or on 3D statistical images of the activations (image-based meta-analysis, IBMA) (Salimi-Khorshidi et al., 2009) (Gorgolewski et al., 2015). The BrainMap project represents a pioneering effort to establish standardized methods for spatial normalization, referencing to common coordinates, and linking coordinates and images to associated literature (Fox and Lancaster, 2002) (Fox et al., 2014). These and other fMRI data-sharing efforts (Maumet et al., 2016) (Reid et al., 2016) have resulted in an explosion of fMRI meta-analyses (Costafreda et al., 2008) (Hart et al., 2013) (Barquero et al., 2014) that have increased understanding of basic brain function and have enabled the development of tools and biomarkers for diagnosis of and evaluation of treatments for mental illnesses.

Poldrack et al. (Poldrack et al., 2017) lays out a comprehensive strategy for transparent and reproducible fMRI imaging and makes a strong case for the need for standardized automated processing, benchmarks, and large-scale sharing of raw data as well as statistical maps. The BIDS (Brain Imaging Data Structure) specification (Gorgolewski et al., 2016) represents a significant international effort to develop data standards for raw and processed neuroimaging data and associated metadata. The BIDS standardization and support tools for fMRI data are relatively mature. In contrast to meta-analysis, *mega-analysis* involves the pooling and joint analysis of raw data across recordings and/or studies, rather than combining metadata or derived statistics. With the creation of open-access platforms for sharing raw data such as OpenfMRI (Poldrack et al., 2013) and its successor OpenNeuro (Gorgolewski et al., 2017), fMRI *mega-analyses* are now starting to appear.

While OpenNeuro is currently dominated by fMRI data, the recent adoption of BIDS extensions for MEG (magnetoencephalography) and EEG (electroencephalography) imaging (Niso et al., 2018) has greatly increased interest in making EEG data publicly available on OpenNeuro and other platforms. Unfortunately, there is no EEG/MEG equivalent for the standardized statistical maps that have powered successful fMRI meta-analyses. Furthermore, although several general guidelines for best practices in EEG/MEG data acquisition and preprocessing have recently appeared (Gross et al., 2013) (Keil et al., 2014) (Pernet et al., 2018), MEG/EEG preprocessing has not been standardized and is only starting to be systematically benchmarked (Pedroni et al., 2018).

Central to mega-analysis, which combines raw data across recordings and studies, are the issues of how EEG data should be normalized to improve comparability of signals across studies. This paper examines channel-level and source-level spatial, temporal, and spectral signal properties across 18 studies performed at six different experimental sites in an effort to provide preliminary guidance on these issues. A companion paper (Bigdely-Shamlo et al., 2018) applies a more detailed mega-analysis with hierarchical statistical modeling to assess commonalities in event-related temporal and spectral features across the same corpus.

The corpus includes several groups of studies that use the same general paradigm but differ in protocol details. For example, five of the studies are based on the RSVP (Rapid Serial Visual Presentation) paradigm (Bigdely-Shamlo et al., 2008; Sajda et al., 2003), but have different types of images, rates of presentation, and requirements for manual response to mark target detection. Other studies use tasks such as lane-keeping with vehicle perturbations in a driving simulation. This corpus, which reflects the interests of the participating research groups, is indicative of the types of studies and heterogeneity that are likely to become available on public repositories where control variables across studies may not be comparable, even when multi-factor designs are utilized within individual studies. Further, there is no guarantee that subject IDs used within a study are consistent across studies performed at a given institution, thus systematic accounting of cross-subject variability is difficult in consolidated analyses. However, even in the unstructured environments likely to be encountered in public repositories, there are universal elements that can be explored and exploited.

Over the last decade, there has been an increased emphasis on the use of multi-level statistical models (e.g., hierarchical linear models or mixed effects models) in EEG data analysis to jointly analyze variability both within and between subjects, groups, and studies (Pernet et al., 2011) (Stefan J. Kiebel and Friston, 2004) (Stefan J Kiebel and Friston, 2004). Explicitly modeling recording- and study-specific variability in a partial-pooling analysis can help address issues with Type I error rate inflation associated with classical full-pooling analyses. In EEG analysis, such multi-level models are typically applied to a structured corpus of event-related data. In this paper, we study the general characteristics of the data using full pooling analyses. We defer to our companion paper (Bigdely-Shamlo et al., 2018) a more detailed multi-level statistical analysis in which we characterize the effects of various experimental factors on event-related temporal and spectral features, while explicitly modeling recording-specific variability using a two-level hierarchical linear model.

This paper is organized into two main themes in which we examine commonalities in channel-level and source-level spatial, temporal, and spectral properties across studies. The Methods section briefly describes the experimental data, provides an overview of the automated processing pipeline used to perform the analysis, and introduces a class of robust measures to explore normalization and statistical properties of EEG across heterogeneous studies. The Results section follows a similar organization with a presentation of results on statistical differences and similarities of EEG signals in both channel (section 3.1 Channel analysis) and source (section 3.2 Source analysis) space. The Discussion section offers a perspective on the analysis of heterogeneous datasets and directions for future mega-analyses. Together, this work provides a broad insight into the sources of variability contained within a large, heterogeneous dataset of EEG recordings and how these factors can be addressed to reveal consistent differences in power spectral amplitude across frequency bands and brain regions.

## 2 Methods

### 2.1 Experimental data

As summarized in Table 1, this paper uses data from 18 studies conducted at six sites associated with four institutions: the Army Research Laboratory (ARL), the National Chiao Tung University (NCTU), the Netherlands Organization for Applied Scientific Research (TNO), and the University of California at San Diego (UCSD). All studies were conducted with voluntary, fully-informed subject consent and approved by the Institutional Review Boards of the respective institutions. More detailed information about the studies with explanations of acronyms and relevant citations appear in Appendix A.

**Table 1:**
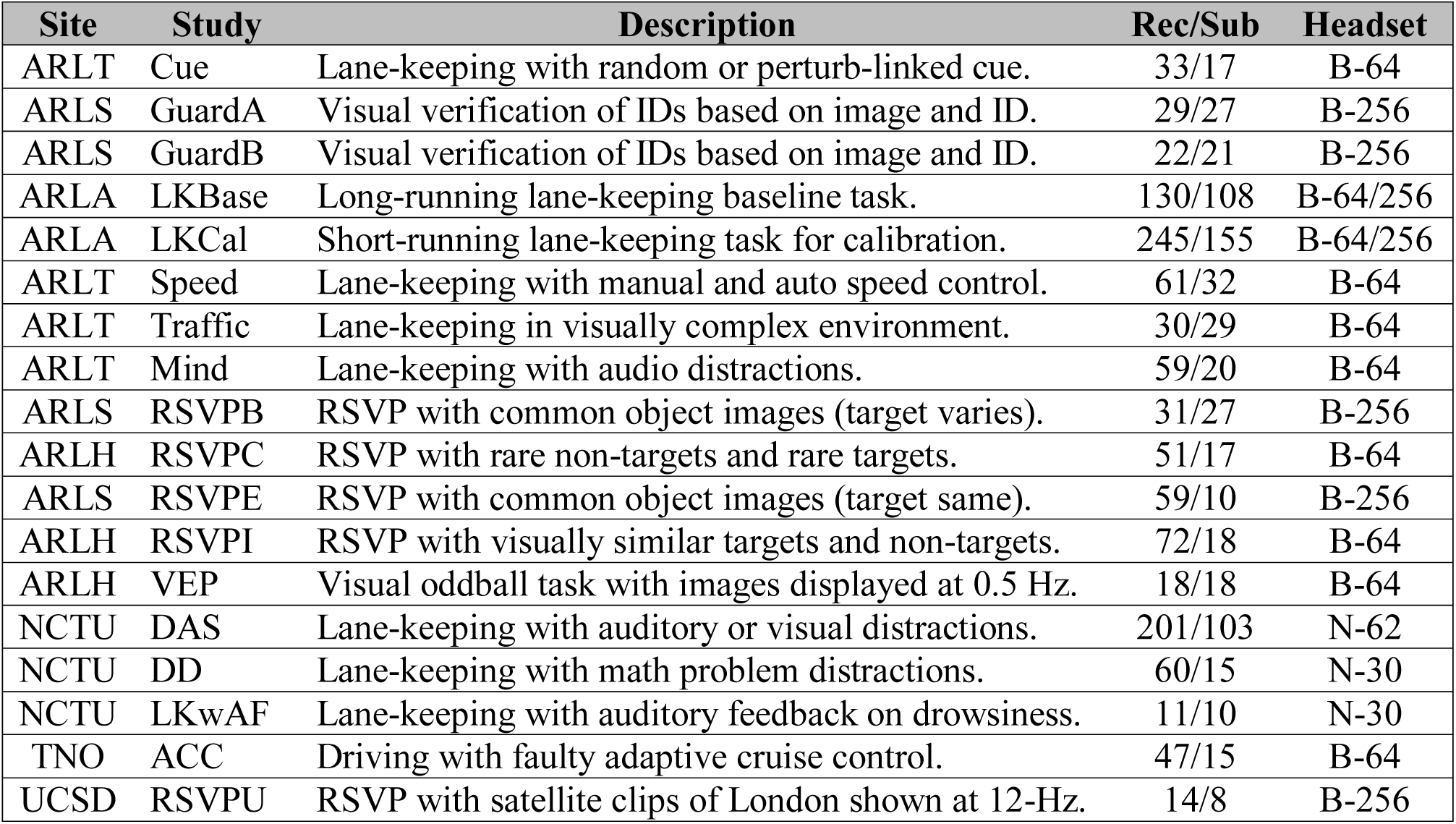
Summary of datasets used. The Rec/Sub column gives the number of recordings and the number of unique subjects within a study. Subjects may overlap across studies from the same site. Headset (B = BioSemi, N = Neuroscan) specifies headset type and number of channels.

| Site | Study | Description | Rec/Sub | Headset |
| --- | --- | --- | --- | --- |
| ARLT | Cue | Lane-keeping with random or perturb-linked cue. | 33/17 | B-64 |
| ARLS | GuardA | Visual verification of IDs based on image and ID. | 29/27 | B-256 |
| ARLS | GuardB | Visual verification of IDs based on image and ID. | 22/21 | B-256 |
| ARLA | LKBase | Long-running lane-keeping baseline task. | 130/108 | B-64/256 |
| ARLA | LKCal | Short-running lane-keeping task for calibration. | 245/155 | B-64/256 |
| ARLT | Speed | Lane-keeping with manual and auto speed control. | 61/32 | B-64 |
| ARLT | Traffic | Lane-keeping in visually complex environment. | 30/29 | B-64 |
| ARLT | Mind | Lane-keeping with audio distractions. | 59/20 | B-64 |
| ARLS | RSVPB | RSVP with common object images (target varies). | 31/27 | B-256 |
| ARLH | RSVPC | RSVP with rare non-targets and rare targets. | 51/17 | B-64 |
| ARLS | RSVPE | RSVP with common object images (target same). | 59/10 | B-256 |
| ARLH | RSVPI | RSVP with visually similar targets and non-targets. | 72/18 | B-64 |
| ARLH | VEP | Visual oddball task with images displayed at 0.5 Hz. | 18/18 | B-64 |
| NCTU | DAS | Lane-keeping with auditory or visual distractions. | 201/103 | N-62 |
| NCTU | DD | Lane-keeping with math problem distractions. | 60/15 | N-30 |
| NCTU | LKwAF | Lane-keeping with auditory feedback on drowsiness. | 11/10 | N-30 |
| TNO | ACC | Driving with faulty adaptive cruise control. | 47/15 | B-64 |
| UCSD | RSVPU | RSVP with satellite clips of London shown at 12-Hz. | 14/8 | B-256 |

The studies included variations of tasks that fall into two general categories: visual target detection and driving with/without motion platform, traffic, speed control, distraction, auditory feedback, or adaptive cruise control. The visual target detection tasks included simulated guard duty with ID checks (GUARDA and GUARDB), rapid serial visual presentation (RSVPB, RSVPC, RSVPE, RSVPI, RSVPU), and visual oddball (VEP). The other tasks were all based on variations of lane-keeping in a driving simulator, with the exception of ACC, which involved driving in an actual vehicle on a test track. In all, the corpus includes 1,173 EEG recordings running 633 hours. The NCTU, TNO and UCSD experiments were all performed at a single site at the respective institutions, while the ARL experiments were performed at three different sites. ARLH designates experiments performed at the Human Research and Engineering Directorate at Army Research Laboratory (Aberdeen, MD), ARLS designates experiments performed at SAIC Laboratories (Louisville, CO), and ARLT designates experiments performed at Teledyne Corporation (Durham, NC). ARLA designates tasks (LKBase and LKCal) that were performed at all three sites using identical experimental setups.

To facilitate consistent large-scale downstream processing, the entire corpus was organized in a containerized form using ESS (EEG Study Schema) (Bigdely-Shamlo et al., 2016b) (*BigEEG Workflow*, 2018). The individual EEG recordings were converted to EEGLAB set format (Delorme et al., 2011), and the study-specific event codes were mapped to Hierarchical Event Descriptor (*HED*) strings (Bigdely-Shamlo et al., 2016a).

### 2.2 Data organization and preprocessing

Data cleaning was fully-automated and proceeded in several stages. (See Appendix B for a summary graphic of the preprocessing procedure.) We used *PREP* (Bigdely-Shamlo et al., 2015), an automated open-source tool that we developed, to perform robust average referencing and interpolation of bad channels. After removing line noise from each data recording, *PREP* identifies and interpolates noisy channels to form a robust average reference. We also removed non-EEG channels from each recording and assigned 10-20 channel labels to channels that did not have standard labels (e.g., 256-channel BIOSEMI caps) based on channel distances to standard 10-20 locations (Klem et al., 1999) (Jurcak et al., 2007). After selecting a maximum of 64 channels based on the 64-channel 10-20 configuration for this analysis, we resampled the data at 128 Hz using EEGLAB *pop_resample()*. Since all 18 studies used in this paper were acquired at rates higher than 128 Hz, resampling is actually down-sampling. We applied a 1690-point high-pass filter at 1 Hz with the *pop_eegfiltnew()* function of EEGLAB to remove low-frequency drift.

We used a combination of independent component analysis (ICA) and regression of residual blink activity to remove eye activity from the corpus. After performing Infomax ICA (Bell and Sejnowski, 1995) using the portion of the data without outlier amplitudes as described in Mullen et al. (Mullen et al., 2015), we applied a modified version of *EyeCatch* (Bigdely-Shamlo et al., 2013a) to identify independent components (ICs) associated with eye activity. *EyeCatch*, which contains a database of 3,453 manually identified eye-related ICA scalp maps, flags “eye” ICs as those that either have a similar topography (correlation > 0.94) to at least one scalp map from the database or are moderately similar (correlation > 0.85) and have a power ratio for at least 100 for at least 1% of data frames. To compute the power ratio, *EyeCatch* performs a continuous wavelet transform (Lilly and Olhede, 2012; Mallet, 2008) using the MATLAB *cwt()* function and calculates the power in the range [1, 3] Hz divided by the power in the range (3, 15] Hz at each time point. We then removed the subspace spanned by the ICs identified by *EyeCatch* as containing eye activity from the channel data.

We also applied *BLINKER* (Kleifges et al., 2017) to channel data to identify latencies associated with different phases of eye blinks prior to eye artifact removal. *BLINKER* inserts five events into the EEG.event structure (*Left base*, *Left zero*, *Blink peak*, *Right zero*, *Right base*) for each blink. *BLINKER* also extracts a continuous “blink” signal that follows the blink-induced EEG activity between each pair of [*Left zero*, *Right zero*] markers and is zero outside these intervals. We applied temporal overlap regression using a method similar to that proposed by Kristensen et al. (Kristensen et al., 2017) to regress out both the effects of blink events and the continuous blink signal produced by *BLINKER*, assuming blinks are fixed patterns in an interval [−1, 1] seconds time-locked to the *Blink peak* events.

### 2.3 Channel analysis

The goal of channel analysis is to assess the statistics and variability of channel data across studies. Several analysis steps normalize the data by removing the median and dividing by the robust standard deviation for each channel. We refer to this normalization as a “robust z-score” data transformation. The robust standard deviation is defined as 1.4826 times the median absolute deviation from the median (*MAD*). In this paper, we refer to the robust standard deviation of an EEG channel as the “robust channel amplitude.”

#### 2.3.1 Variation of channel amplitudes across scalp positions and recordings

To investigate whether EEG signals have consistently different amplitudes across channel locations and recordings, we developed a compact representation of the amplitude of each recording by a 26×1 positive vector called the “recording amplitude vector”. We calculated the recording amplitude vector by first selecting the following subset of 26 channels common across all of the recordings: Fp1, Fp2, F3, Fz, F4, F7, F8, FC3, FCz, FC4, FT7, FT8, C3, Cz, C4, TP7, TP8, CP3, CPz, CP4, P3, Pz, P4, O1, Oz, and O2. We then calculated the robust standard deviation of each of these channels after low-pass filtering channel signals at 20 Hz. To perform analysis across *R* recordings, we formed the “amplitude matrix”, a 26×*R* matrix of these column vectors. (Note, this low-pass filtering and channel down-selection was only used for the purpose of channel analysis and does not apply to the calculation of sources.)

We visualized the overall dependence of robust amplitude on channel position by applying the median function to the rows of the amplitude matrix and then plotting the resulting 26×1 vector using the EEGLAB *topoplot()* function. To visualize potential dependencies of the amplitude vectors on study identity, we applied *t-SNE* (t-distributed stochastic neighbor embedding) (Van Der Maaten and Hinton, 2008) (Van Der Maaten, 2014) to project the columns (recordings) of the amplitude matrix to a 2-D space. We used the MATLAB *tsne()* function with a perplexity parameter of 20 for the visualizations displayed in this paper.

#### 2.3.2 Characterizing variation by a recording-specific constant

A known difficulty with joint analysis of EEG recordings is that the variability introduced by headset technology and sensor placement as well as subject hair, scalp and skull characteristics causes statistical properties of data to vary across recordings. A simple approach for addressing this problem is to model the variability by a recording-specific scaling factor and to “normalize” each recording by dividing its channel data by a recording-specific constant. We investigated several different ways of computing the recording-specific normalization factor including mean, Huber mean (Huber, 1964) (Analytical Methods Committee, 2001), and the Euclidean (*L*^2^) norm of the recording amplitude vector. We also looked at measures based directly on the channel signal rather than the recording amplitude vector, but these methods provided worse comparability and are not reported here. While all of these methods capture the central tendency of recording amplitude, the mean and Euclidean norm are sensitive to outliers. The Huber mean is an iterative technique for robust approximating the mean in the presence of outliers.

The normalization factors were computed column-wise on the amplitude matrix (across channels) as summarized in Figure 1 for the Huber mean. Regardless of the normalization method, the resulting amplitude matrix was divided column-wise by the 1×*R* vector of resulting recording normalization factors to produce a normalized amplitude matrix for each normalization method.

**Figure 1.**
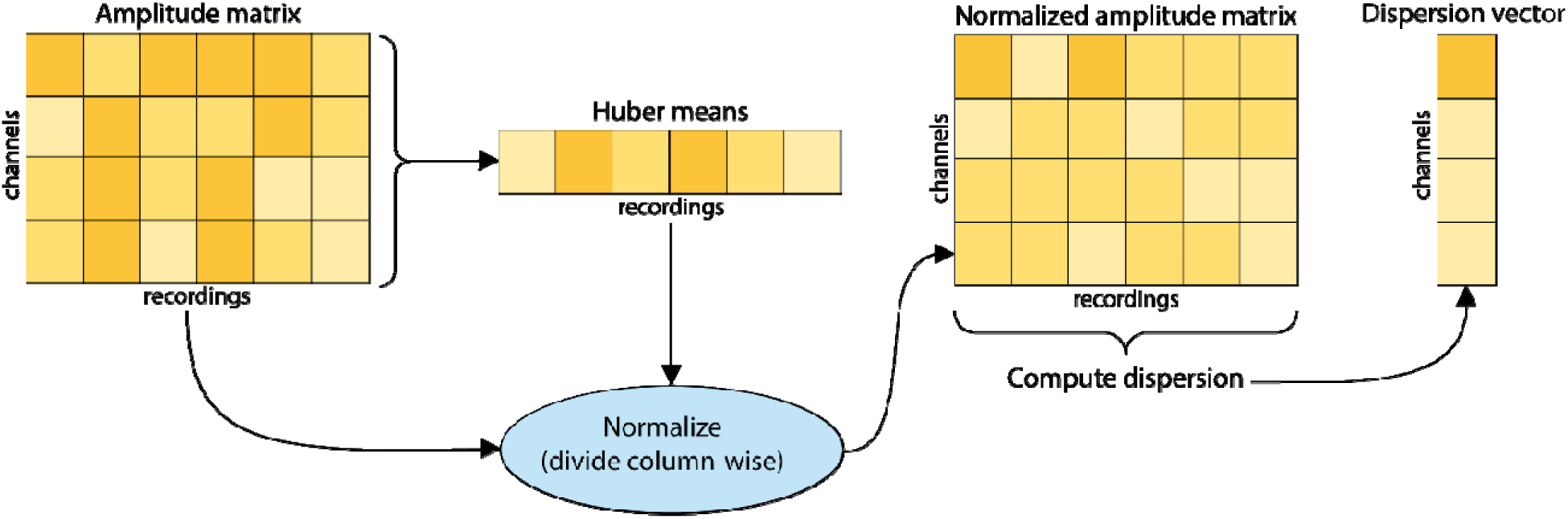
Steps for computing the normalized amplitude matrix and the channel dispersion vector for Huber mean normalization. A similar procedure was used for the other normalization methods.

We performed several tests to assess whether normalizing each recording by a recording-specific constant could improve statistical comparability across recordings from different studies. To characterize comparability of channel amplitudes across a data corpus, we computed the “dispersion vector” by taking the robust standard deviation of each row (across recordings in the corpus) of an amplitude matrix and dividing this vector by the row median (across recordings) to obtain a 26×1 channel dispersion vector representing the corpus variability for each method. We also computed median dispersion across all channels for each channel dispersion vector to find the normalization method that was most effective in reducing overall amplitude dispersion across recordings.

Dispersion vectors for a corpus can be computed at various stages of preprocessing to provide a compact representation of how different processing steps affect channel statistics. We obtained a visual representation of these effects using the EEGLAB *topoplot()* to plot channel dispersion vectors corresponding to different steps and/or preprocessing methods. In order to find out whether the difference in median dispersion was statistically significant for different normalization methods, we applied the pairwise Wilcoxon signed rank test (using MATLAB *signrank()* function) between dispersion vectors calculated using Huber mean (which is the method with the smallest median dispersion) and the dispersion vectors calculated using other normalization methods.

To investigate how much of the channel amplitude variability could be explained by a recording-specific constant scaling of all channel amplitudes in each recording, we plotted channel-pair amplitudes across all recordings and computed adjusted *R*^2^ values for linear fits of data points corresponding to each channel-pair amplitude separately. Significance tests for non-zero slopes of the linear fits were also performed. The same processing steps on channel-pair amplitudes were then repeated after normalizing channel amplitudes in each recording using the recording-specific constant based on Huber mean.

#### 2.3.3 Variation of channel baseline amplitude across scalp positions and frequencies

To investigate whether EEG signals at different frequencies have consistently different amplitudes across channel locations, we analyzed continuous EEG after removing eye activity and normalizing by the Huber recording-specific constant. We selected the same subset of 26 channels used in the investigation of channel signal amplitudes and calculated the continuous wavelet transform (using the MATLAB *cwt()* function) for each channel to obtain a time-varying amplitude spectrogram (square root of the power of 50 frequencies logarithmically sampled between 2 and 40 Hz). The baseline spectral amplitude vector for each channel is a 50 × 1 vector of the median over time of the amplitudes at each of the 50 specified frequencies in the channel spectrogram. To focus on the relative amplitudes of different frequencies in each channel, we normalized each of these baseline spectral amplitude vectors to have Euclidean norm of one (1). We refer to the result as the “normalized channel spectrogram”.

In order to investigate the statistical significance of the observed differences in channel amplitude at each frequency, we applied Wilcoxon signed rank tests (MATLAB *signrank()* function) on the robust z-scores of the channel spectral amplitudes for each (frequency, recording) pair. The null hypothesis of these statistical tests was that for each (frequency, recording) pair, each channel spectral amplitude could be modeled by a constant plus a random value drawn from a zero-median distribution.

#### 2.3.4 Variation of channel covariance across studies

To investigate systematic variations of channel covariance across headsets, paradigms, and other study details, we calculated a single covariance matrix for each recording using the same subset of 26 channels used for the investigation of channel signal amplitudes. We used the same segments of the data that were used to compute Infomax ICA in order to exclude sections with abnormally high power due to artifacts. We plotted the two-dimensional *t-SNE* projections of distances between recording covariance matrices using an EEG headset color scheme to visualize the systematic variations in covariance across recordings with respect to headset. We used the Riemannian distance metric recommended by Förstner and Moonen (Förstner and Moonen, 2003) to compare recording covariance matrices **A** and **B**:

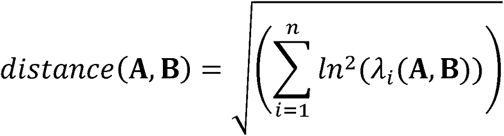

Here *n* is the number of channels, and the λ_*i*_ are generalized eigenvalues. For comparison, we also applied *t-SNE* to the recording channel correlation matrix and to the channel standard deviation vectors (computed as the square root of the variance rather than as the robust standard deviation used in the amplitude analysis). We examined variations with and without recording-specific Huber normalization with both Riemannian distance and Euclidean distance.

We also applied representational similarity analysis (RSA) as proposed by Kriegeskorte et al. (Kriegeskorte et al., 2008) to determine whether similarities in the patterns observed in the *t-SNE* visualizations were statistically significant. Representational analysis determines whether pairwise sample differences associated with two aspects of the data are more correlated than would be expected from chance. The significance tests are performed by random permutation testing. The aspects of the data used here are similarity of covariance matrices of two recordings and whether or not the recordings are from the same headset. In other words, this analysis determines whether covariance matrices from recordings of the same headset type are significantly more correlated than would be expected from chance. More specifically, we used Riemannian distance (converted to a similarity measure) between the covariance matrices of recordings *i* and *j*, respectively, as one measure. The second representation was the headset membership matrix *M*_*ij*_, which has an entry of 1 if recording *i* and recording *j* are from the same headset type and 0 if recording *i* and recording *j* are not from the same headset type.

### 2.4 Source analysis

Our evaluation of brain source signal variation uses sources derived from ICA, a widely-used EEG source analysis technique. We used the ICA components (ICs) computed during preprocessing and not identified as eye activity as the starting point for the analysis.

#### 2.4.1 Variation of brain sources across space and frequency

We inferred equivalent dipole locations for maximally-independent sources of EEG activity by applying the EEGLAB *dipfit* plugin to scalp maps of the ICs not identified as eye artifacts during preprocessing. To further select ICs most likely to be associated with brain sources, we only considered dipoles that matched all the following criteria: (a) the scalp map explained by the dipole accounted for at least 85% of the variance, (b) the EEGLAB *MARA* plugin (Winkler et al., 2014, 2011a) did not identify the component scalp map as an artifact (e.g., muscle or eye), and (c) the dipole was located inside the brain volume, as identified by the *ft_sourcedepth()* function of the EEGLAB *fieldtrip*_*lite* plugin (Oostenveld et al., 2011). Note that each dipole is associated with exactly one recording.

As with channel space analysis, we calculated the continuous wavelet transform (using the MATLAB *cwt()* function) of the continuous time course of the IC activation associated with each dipole to obtain a time-varying amplitude spectrogram (square root of power of 50 frequencies logarithmically sampled between 2 and 40 Hz). The baseline spectral amplitude vector for each IC activation is defined as the 50 × 1 vector of the median over time of the amplitudes at each of the 50 specified frequencies in the IC spectrogram. To focus on the relative amplitudes of different frequencies and to facilitate cross-recording comparison, we scaled each spectrogram so that its overall baseline spectral amplitude vector had Euclidean norm one. We refer to the result as the “normalized spectrogram” for an IC.

To investigate the differences in normalized spectrograms, we converted these values to relative deviations from means, computed at each frequency across all ICs. We define the spectral amplitude deviation value at frequency for the *i*^th^ IC as:

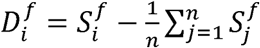

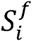 is the normalized spectrogram value at frequency *f* for IC *i* and *n* is the number of ICs. We then used Measure Projection Analysis (*MPA*) to analyze and visualize the spatial distribution of these deviations for different frequency bands (Bigdely-Shamlo et al., 2013b). Using *MPA*, we computed the spatial Gaussian weighted averages (12 mm standard deviation) of IC spectral amplitude deviations for all brain voxels. Averages were also computed for regions of interest (ROIs) selected from the AAL brain atlas (Tzourio-Mazoyer et al., 2002).

#### 2.4.2 Background spectral power characteristics in different brain areas

The power spectrum of electrophysiological signals at the channel or source level can be modeled as a combination of periodic and aperiodic components, reflecting narrowband oscillations (spectral “peaks”) and approximately 1/*f* “background” activity, respectively (Onton and Makeig, 2006) (Miller et al., 2009). While the properties of periodic/oscillatory activity in various frequency bands have been extensively studied and linked to an array of cognitive and behavioral states and neurological disorders, the aperiodic component of the power spectrum has received less attention.

In order to characterize the spatial distribution of the aperiodic spectrum, we computed the Huber mean power spectra across all time points for each IC. We used the Huber mean, instead of median, in order to more closely estimate mean power while ignoring outliers caused by artifacts. We then used the *FOOOF* python toolbox (Haller et al., 2018) (*fooof*, 2018) to model the power spectrum, *P*, as a linear combination of aperiodic background activity, *L*, and a weighted sum of *N* Gaussian functions, *G*_*n*_, each representing a periodic component (i.e., a “peak” in the spectrum):

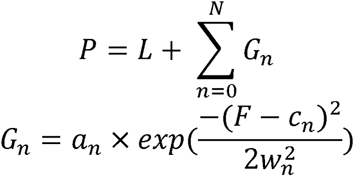

Each Gaussian component, *G*_*n*_, is characterized by its amplitude, *a*_*n*_, its spectral position *c*_*n*_, and RMS bandwidth, *w*_*n*_. *F* is a vector of frequencies. The aperiodic signal *L* is modelled as an exponential function in semilog-power space (linear frequencies and log power values) as:

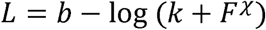

where *b* is the broadband offset, *χ* is the aperiodic slope, and is the “knee” parameter that determines the inflection point of the aperiodic exponential function (Miller et al., 2009).

We set the maximum number of Gaussians to 10 and their width to be between 3 and 20 Hz. This provided *FOOOF* with ample degrees of freedom to model the spectrum and to separate the periodic and aperiodic activity. After fitting each dipole IC power spectrum using *FOOOF*, we removed ICs with a goodness of fit (*R*^2^) less than 0.95. We then computed the average slope, 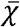, for all ICs and the deviation of each IC’s slope from this average: 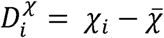.

Similar to the analysis of the spatial distribution of baseline “periodic” amplitude deviations, we analyzed the spatial distribution of aperiodic slope deviations using Measure Projection Analysis (*MPA*), computing the spatial Gaussian weighted averages (12 mm standard deviation) of these values for all brain voxels (Bigdely-Shamlo et al., 2013b).

## 3 Results

### 3.1 Channel analysis

#### 3.1.1 Division by a recording-specific constant improves recording and study comparability

To investigate whether dividing each EEG recording by a single recording-specific number improves the comparability across recordings, we looked at channel amplitude dispersion across studies for different normalization approaches and at different stages in processing. Figure 2 shows the effect of different types of processing on channel amplitude dispersion.

**Figure 2.**
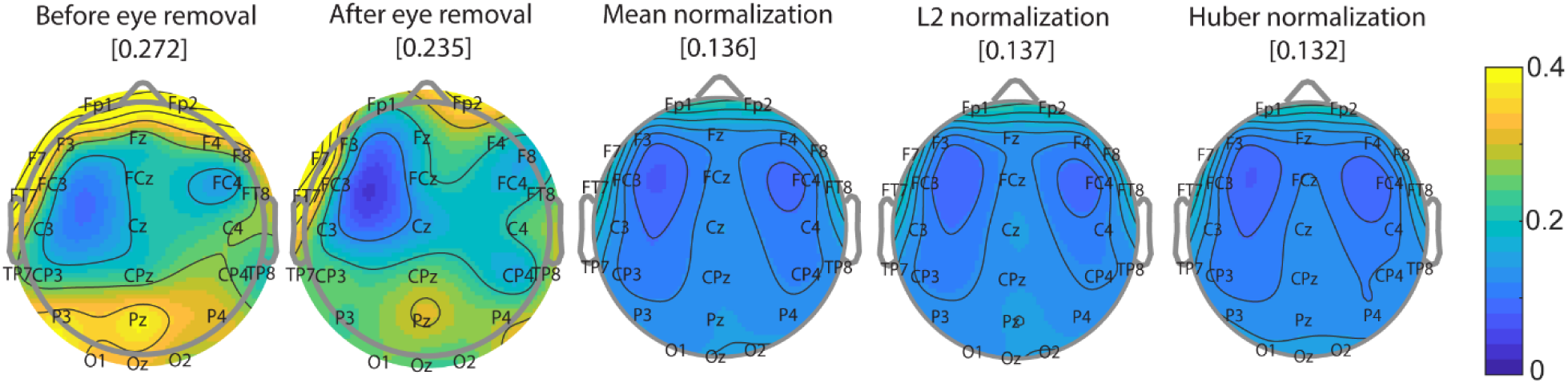
Effect of preprocessing and channel normalization on channel amplitude dispersion. Each scalp map shows the median over all recordings in all studies of the channel dispersion for each channel. The bracketed number over each scalp map is the overall median dispersion corresponding to the scalp map. Normalization always occurs after eye removal.

Interestingly, although removal of eye artifacts changes the spatial distribution of dispersion, it does not substantially reduce the amount of dispersion. Further, the maximum dispersion before eye artifacts are removed does not occur in the frontal channels. Note that dispersion is a dimensionless quantity and scaling a recording by a constant does not change the dispersion. Normalization using any of the three methods (mean, *L*^2^, or Huber) significantly reduces the channel dispersion in comparison with non-normalized recordings. Huber mean and mean are essentially the same for datasets without outliers, but Huber mean produces a slightly lower dispersion when there are outliers. The spatial distribution of dispersion is similar for all three normalization methods.

Figure 3 shows the distribution of channel dispersions segregated by study before removal of eye artifacts (black box plots), after removal of eye artifacts (blue box plots), and after eye artifact removal followed by Huber normalization (green box plots). Huber normalization significantly reduces the median dispersion across all studies. Normalization also reduces the number of channel dispersion outliers for most studies. Note that LKBase and LKCal were basic lane-keeping studies that included recordings acquired at three different experimental sites (HRED Army Research Laboratory in Aberdeen, MD, Teledyne Laboratories in Durham, NC, and SAIC Laboratories in Louisville, CO) using two different headset types. The channel dispersions across these tasks were well clustered and did not show a particular dependence on site location.

**Figure 3.**
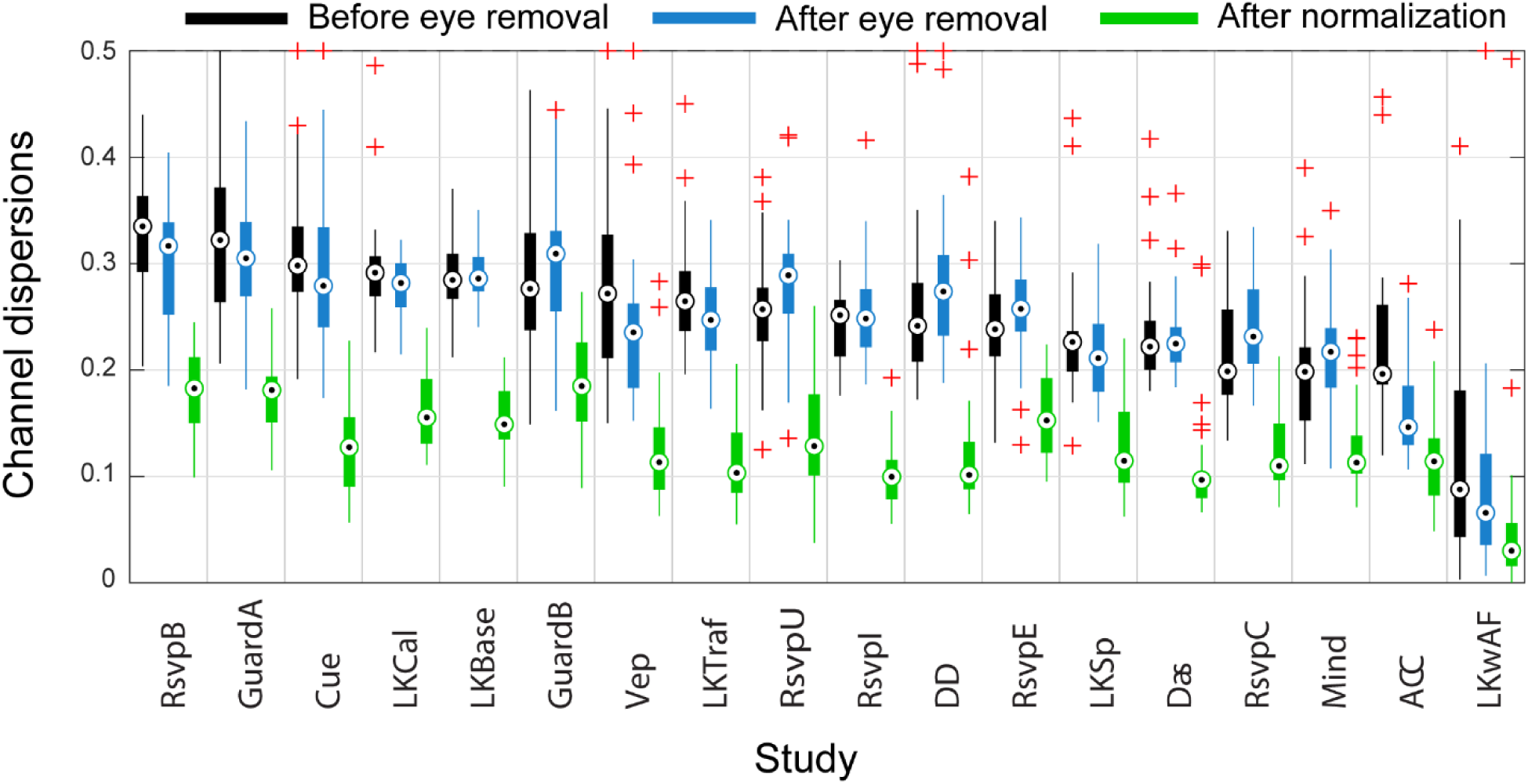
Channel dispersions segregated by study before eye artifact removal (black boxplots), after eye artifact removal (blue boxplots), and after Huber normalization (green boxplots). Circles mark the medians and plus signs mark boxplot outliers. Studies are sorted by the median channel dispersion before eye removal.

#### 3.1.2 Some channels consistently have a higher amplitude than others

Both removal of eye artifacts and scaling by a recording-specific constant affect the relative distributions of channel amplitudes as illustrated in Figure 4. The first row of Figure 4 displays results computed on the composite channel amplitude matrix, corresponding to continuous EEG data after 128 Hz down-sampling, but before removal of eye artifacts. The middle row of Figure 4 displays results computed on the composite channel amplitude matrix after removal of eye artifacts. The bottom row of Figure 4 displays the results based on the composite channel amplitude matrix after eye artifacts have been removed and each recording is scaled by a recording-specific value (Huber mean).

**Figure 4.**
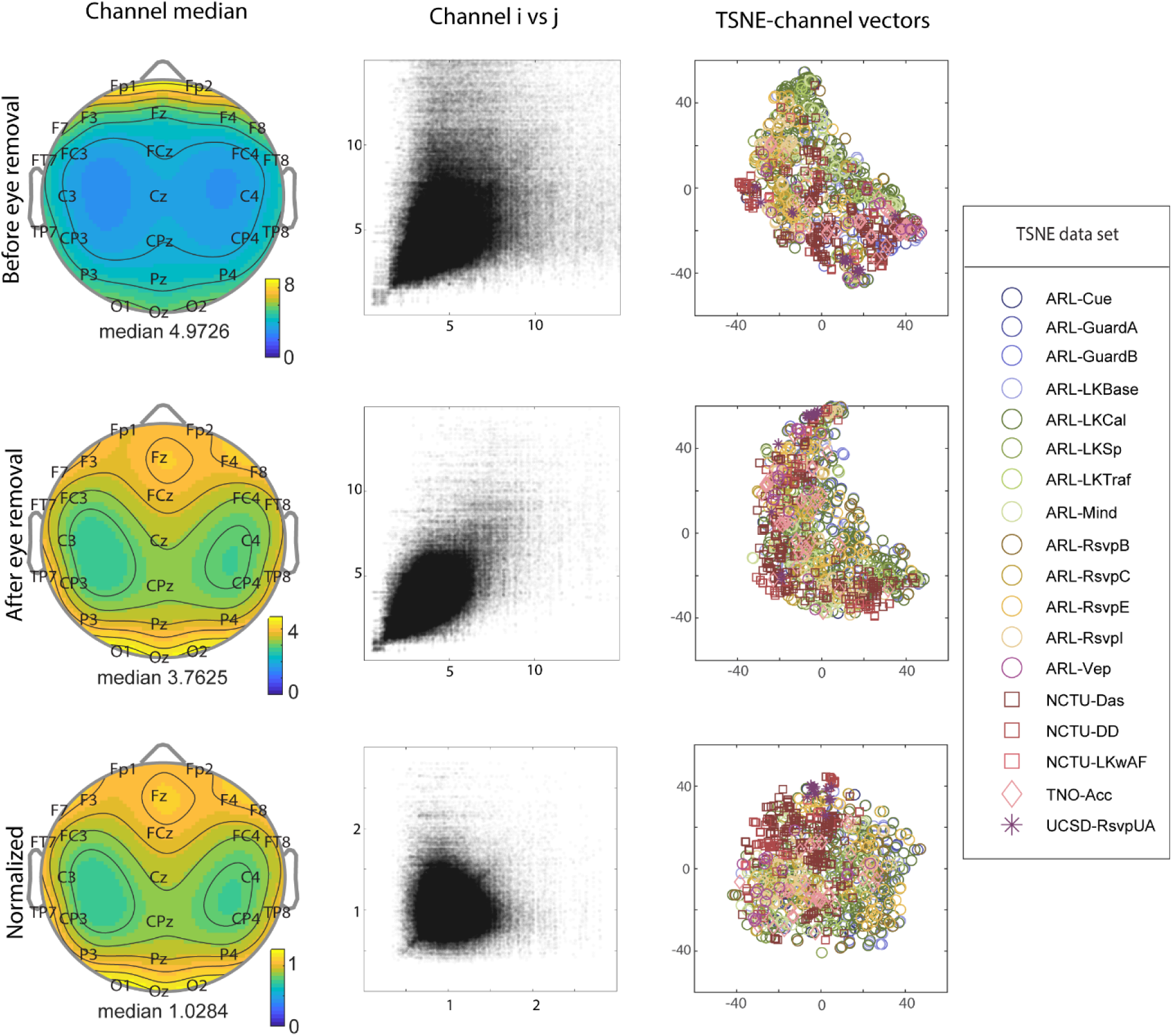
Composite statistics of robust channel amplitudes. Top row uses the composite channel amplitude matrix before eye artifact removal, the middle row uses the composite channel amplitude matrix after eye artifact removal, and the bottom row uses the composite channel amplitude matrix after eye artifact removal and Huber normalization. The first column displays scalp maps of the row medians of the composite channel amplitude matrix at various stages in processing. The middle column plots the robust channel amplitude of channel *i* versus channel *j* (*j* < *i*) in each recording. The last column shows the two-dimensional *t*-*SNE* projection of recording channel amplitude vectors across all studies. The points for each study are depicted using unique colors, and point shape designates the institution (with the three ARL sites combined).

The first column of Figure 4 shows the scalp maps of the row medians of the respective channel amplitude matrices: before eye artifact removal, after eye artifact removal, and after Huber normalization, respectively. The display uses the EEGLAB *topoplot()* function and the 26 common channels listed in the Methods section. The median robust channel amplitude is reduced after eye artifact removal (3.76 versus 4.97), and the spatial distribution of median robust channel amplitude goes from being concentrated in the frontal areas to a two-lobe form with amplitude more evenly distributed but concentrated in the frontal and occipital regions of the scalp. Notice that normalization does not markedly change the spatial distribution of the scalp map, since normalization just corresponds to a reweighting of the points contributing to the median. However, normalization does substantially reduce the overall median robust channel amplitude (to 1.03).

Statistical testing (Wilcoxon signed rank) of robust channel amplitudes for the 26 common channels showed that prior to eye artifact removal all channels had robust amplitudes that differed significantly (*p* < 0.01, *FDR* corrected) from the median of the robust channel amplitude in each recording. After eye artifact removal and after normalization, Fp2 was the only channel that did not differ significantly different from the recording median.

The middle column of Figure 4 plots robust amplitude of channel *i* versus channel *j* (*j* < *i* to eliminate duplicates) in each recording before removal of eye artifacts (top row), after removal of eye artifacts (middle row), and after Huber normalization (bottom row). The top graph is somewhat asymmetric with respect to the 45° line, reflecting the amplitude dominance of the frontal channels. After eye artifacts are removed, the relationship becomes more symmetric, with a visible linear trend, suggesting the existence of an overall recording-level uniform scaling of channel amplitudes. After dividing the channel data by the recording Huber mean normalization factor (an overall robust measure of the recording’s channel amplitude), the linear trend between channel pair amplitudes appears to be essentially eliminated (bottom graph). The median adjusted-R-squared value, resulting from fitting a linear regression models to each group of channel amplitude pairs, is 0.471 before eye artifact removal, 0.488 after eye artifact removal, and 0.014 after normalization. Using data after eye artifact removal and performing linear fits separately for different channel pairs (each fit on amplitudes across recordings), we find that in 100% of these linear fits, the slope factor is nonzero (*p* < 0.01, *FDR* corrected). This linear relationship, which explains about half of the variability in channel pair amplitudes, mostly disappears after Huber mean normalization.

The rightmost column of Figure 4 shows the *t*-*SNE* projections of the recording amplitude vectors into a two-dimensional plane. Although the area occupied by the projections is more circular after normalization, the points are highly overlapping both before and after Huber normalization. The top row, which corresponds to the recordings before removal of eye artifacts shows more segregation by study, emphasizing the importance of consistent eye-artifact removal in cross-study analysis.

#### 3.1.3 Channel covariance exhibits a strong dependence on EEG cap

To understand whether there are systematic variations in channel covariance across studies, we vectorized recording covariance matrices (using the 26 common channels defined in previous sections) and applied *t-SNE* to project these vectors into a two-dimensional plane.

Figure 5 shows the *t*-*SNE* covariance projections plotted with headset type labeled using a unique symbol-color pairing. The N30 (gold) and N62 (red) point groups correspond to recordings from three studies (DAS, DD, LKwAF) performed at NCTU using Neuroscan headsets in two electrode configurations. These points correspond to different tasks and many different subjects. The B256_A point group, which uses a custom cap and an older Biosemi 256-channel headset, corresponds a single study (RSVPU) of 14 recordings from 8 subjects performing an RSVP task. The B256 point group corresponds to data from studies conducted at ARLS using a 256-channel headset. The points in this group represent a number of subjects and several studies (GuardA, GuardB, RSVPB and RSVPE), as well as a subset of LKBase and LKCal recordings. All of these studies were conducted at a single site (ARLS). We also created plots using an individual marker/color type for each study, but did not observe any point segregation by study that was not explained by headset.

**Figure 5.**
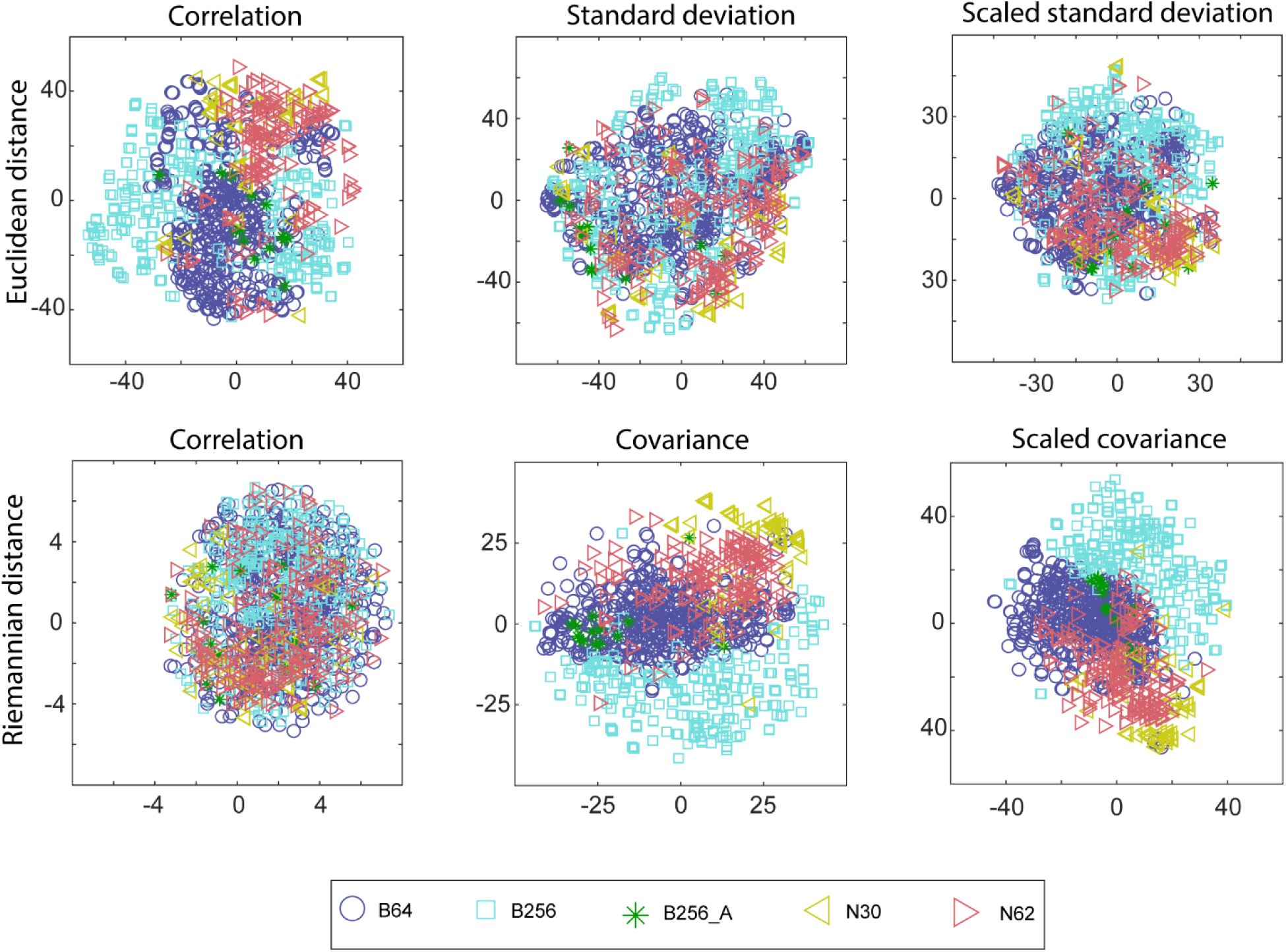
Two-dimensional *t*-*SNE* visualization of selected vectorized recording signal functions. The top row of plots uses Euclidean distance and the bottom row uses Riemannian distance. Points are colored by headset type: B64 designates a Biosemi 64-channel headset, B256 designates a Biosemi 256-channel headset, B256_A designates the UCSD Biosemi 256-channel headset with a custom cap, N30 designates a Neuroscan 30-channel headset, and N62 designates a Neuroscan 62-channel headset.

Visual inspection of the top row of Figure 5, which uses Euclidean distance for the *t-SNE* computation, suggests a possible headset relationship only in the scaled standard deviation. This measure was computed by dividing the standard deviation (a channel amplitude measure) by the recording-specific Huber mean.

Riemannian distance, used in bottom row of Figure 5, is scale-free in the sense that multiplying two covariance matrices by the same constant does not change their distance. The metric is only weakly dependent on scale when two covariance matrices are multiplied by different constants. Among the plots in the bottom row of Figure 5, the covariance and scaled (divided by squared recording-specific Huber mean) covariance plots show distinct clusters, suggesting a strong relationship with headset. The correlation plot, however, does not present any prominent clustering, suggesting a weak or non-existent relationship with headset. We did not include a scaled correlation plot since scaling does not affect correlation, which is intrinsically scale-free.

To quantify the strength and statistical significance of these relationships, we applied Representational Similarity Analysis (RSA) to several different transformations of the recording channel signals, including the vectorized channel correlation, channel covariance, and channel standard deviation. The RSA question we investigated was whether membership of two recordings in the same headset class implied that their signal behavior was more likely to be similar than would be expected if the recordings were assigned to a membership class at random. Table 2 summarizes the RSA statistics corresponding to Figure 5. All of the results of the table, except for the relation between correlations using the Riemannian metric, were significant with p < 0.0005. A higher RSA correlation means that a signal function has a stronger relationship with headset class.

**Table 2:** RSA results for statistical significance of the relationship of several channel signal characteristics (standard deviation, correlation, and covariance) and the membership function of headset type. All results were statistically significant *p* < 0.0005.

| Signal function | Distance metric | Scaling | p-value | RSA correlation |
| --- | --- | --- | --- | --- |
| Correlation | Euclidean | | $\leq 0.0005$ | 0.19 |
| Standard deviation | Euclidean | | $\leq 0.0005$ | 0.04 |
| Standard deviation | Euclidean | Huber mean | $\leq 0.0005$ | 0.08 |
| Correlation | Riemannian |  | 0.1810 | 0.00 |
| Covariance | Riemannian | | $\leq 0.0005$ | 0.29 |
| Covariance | Riemannian | Huber mean | $\leq 0.0005$ | 0.30 |

In this context, the two likely sources of variability affecting channel amplitudes are headset and subjects. A hypothesis supported by the RSA correlation is that scaling of standard deviation by the recording-specific Huber mean removes some of the variability due to subject, thereby strengthening the relative relationship to headset. This is supported by the standard deviation results of Table 2: the RSA correlation increases from 0.04 to 0.08 after normalization, reflecting a stronger relationship to headset.

In Table 2 we also find some evidence for the hypothesis that headsets affect the relative amplitude of channels beyond a uniform scaling effect; the relationship of covariance with headset, using the Riemannian metric, is relatively strong (0.29) and only slightly increased (to 0.30) by scaling. However, this was not the case for channel correlations using the Riemannian metric. Since the only difference between correlation and covariance is the relative channel amplitude, it is likely that headset type primarily affects the relative amplitude of channels (but not their correlation structure). A simple explanation for the headset dependence is the differences in sensor positions associated with the 10-20 mappings, since we chose not to interpolate channel positions to an exact standard. These differences are explored in Appendix C. Other potential sources beyond variation in sensors and their positions is that the pattern of sensor pressures on the scalp exerted by the 256-channel caps may be different from that exerted by caps with lower numbers of sensors. This would explain the relative distances between clusters observed in Figure 5 (scaled covariance): 256-channel headset groups (B256_A and B256), followed by B64, N62 and finally N30 on the corner furthest from all other groups. These systematic differences merit further investigation.

#### 3.1.4 Some channels consistently have higher amplitude at certain frequencies

Statistical testing (Wilcoxon signed rank) also shows that channel amplitudes consistently vary at different frequencies. Figure 6 shows normalized channel amplitudes at different frequencies, each masked by *p* < 0.01 (*FDR* corrected). The amplitude patterns between 4 and 8 Hz resemble the median channel amplitude plots in Figure 4 after eye activity removal, suggesting that overall channel amplitude patterns are highly influenced by amplitude differences in the 4 to 8 Hz range. Note that the channel amplitude vector before eye removal more closely resembles the spatial distribution observed in the 15 to 30 Hz region of Figure 6.

**Figure 6.**
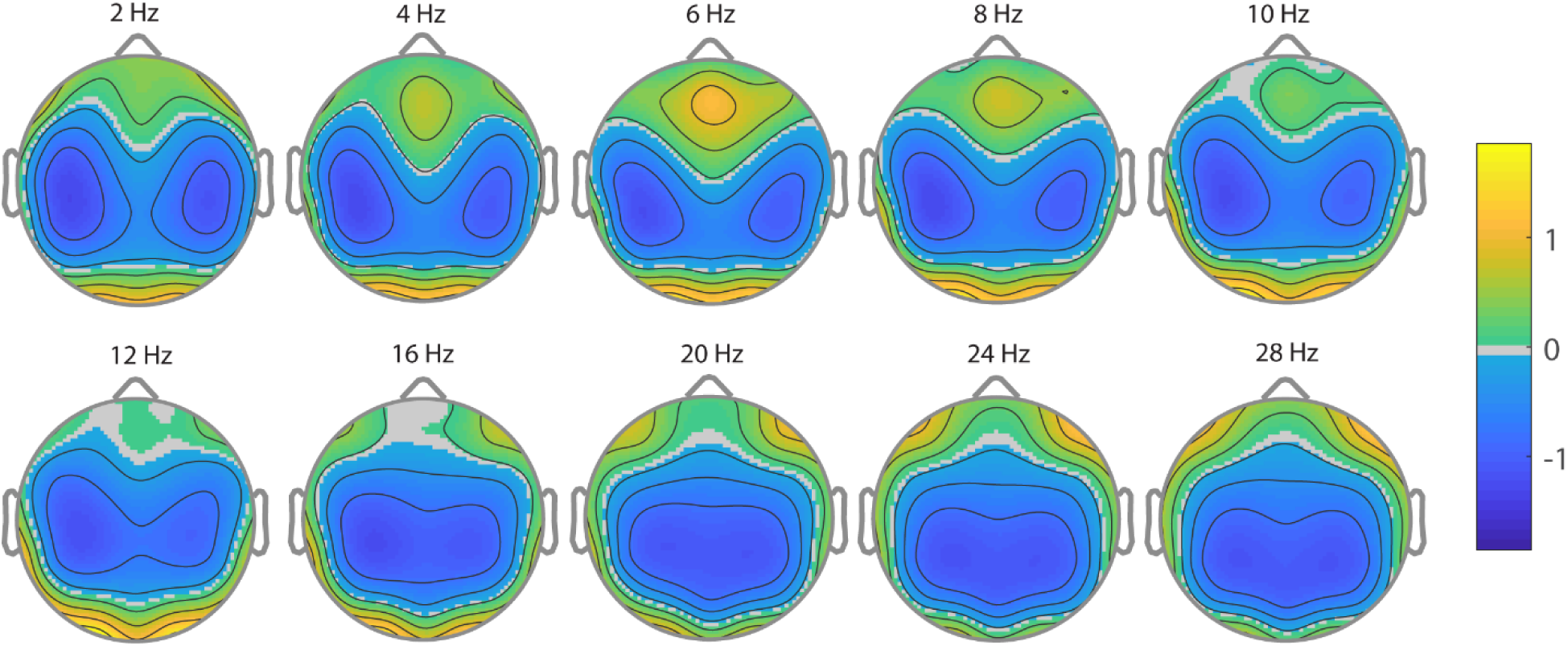
Median normalized channel amplitudes across recordings from all studies at different frequencies, masked by *p* < 0.01, *FDR* corrected after removal of eye artifacts and normalization by a recording-specific constant.

### 3.2 Source analysis

#### 3.2.1 Some brain areas consistently have higher power in certain frequencies

To investigate whether there are systematic variations in amplitude associated with particular frequency bands and brain areas, we applied measure projection analysis (MPA) to the spectrogram deviations calculated from the in-brain IC dipoles (residual variance < 0.15) across the corpus and mapped the results to brain areas using the AAL atlas as shown in Figure 7. There were originally 56,574 dipoles of which 26,175 were in the brain. Applying the requirement of residual dipole variance 0.15 left 11,637 dipoles.

**Figure 7.**
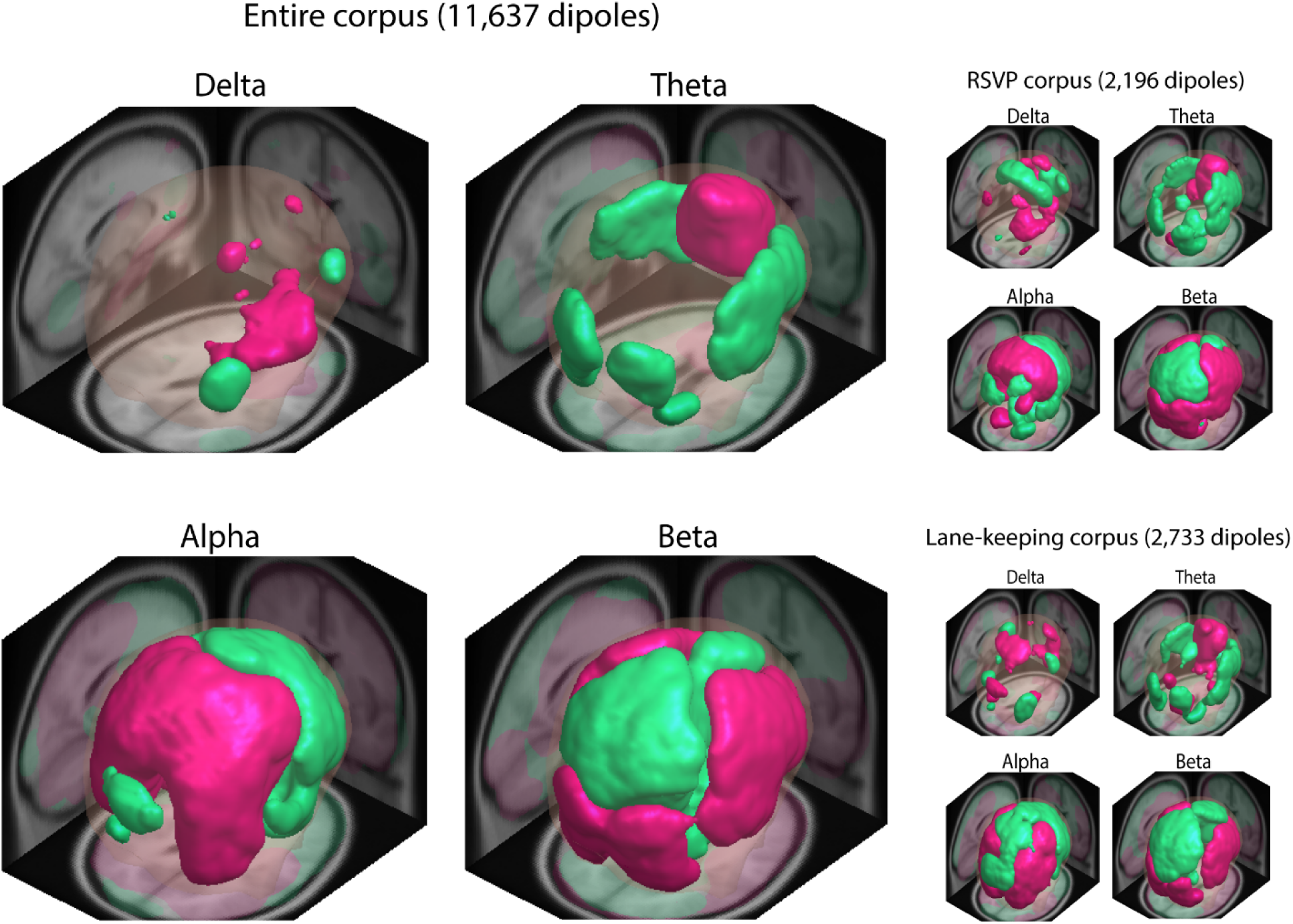
Brain regions with local average (12 mm spatial Gaussian smoothing) spectral amplitude deviation more than 5% (red) and less than −5% (green) for common frequency bands. The left column shows results from the entire corpus, while the right column separately shows the results for two distinct groups within the corpus – one group (*Rsvp*) consists of the five RSVP studies and the other (*Lane-keeping*) consists of data from LKBase and LKCal.

The left column of Figure 7 shows three-dimensional views of brain areas with high (red) or low (green) local spectral amplitude deviation averages in different frequency ranges: delta ([2, 4] Hz), theta ([4, 8] Hz), alpha ([8, 14] Hz), and beta ([14, 31] Hz). Brain areas where the average local amplitude deviation, obtained via 12 mm standard deviation 3D Gaussian smoothing, within the specified frequency band was at least 5% are shown in red. Regions where the amplitude deviation was at most −5% are coded in green.

These 3D views show a clear anterior/posterior dichotomy in alpha amplitudes: posterior brain areas (in red) generally have more alpha amplitude than anterior areas (in green). Beta activity appears to be increased in sensorimotor, middle frontal, and temporal regions. We also note increased activity in inferior occipital and cerebellar regions, which may reflect neck muscle activity (Gramann et al., 2010). Beta activity is reduced along central midline structures, including anterior cingulate, superior frontal, superior parietal, posterior cingulate, and medial occipital regions. We also observe frontal theta activity in anterior cingulate and superior frontal gyrus. Baseline levels were computed by random permutation.

We investigated whether these differences are statistically significant, as shown in Figure 8 for the entire data corpus. The top row of the figure shows transverse brain slides overlaid with dipole density images. The dipole density was computed using Gaussian spatial smoothing with a standard deviation of 12 mm. While dorsal and midline brain regions have a higher dipole density, dipoles are distributed throughout the brain volume

**Figure 8.**
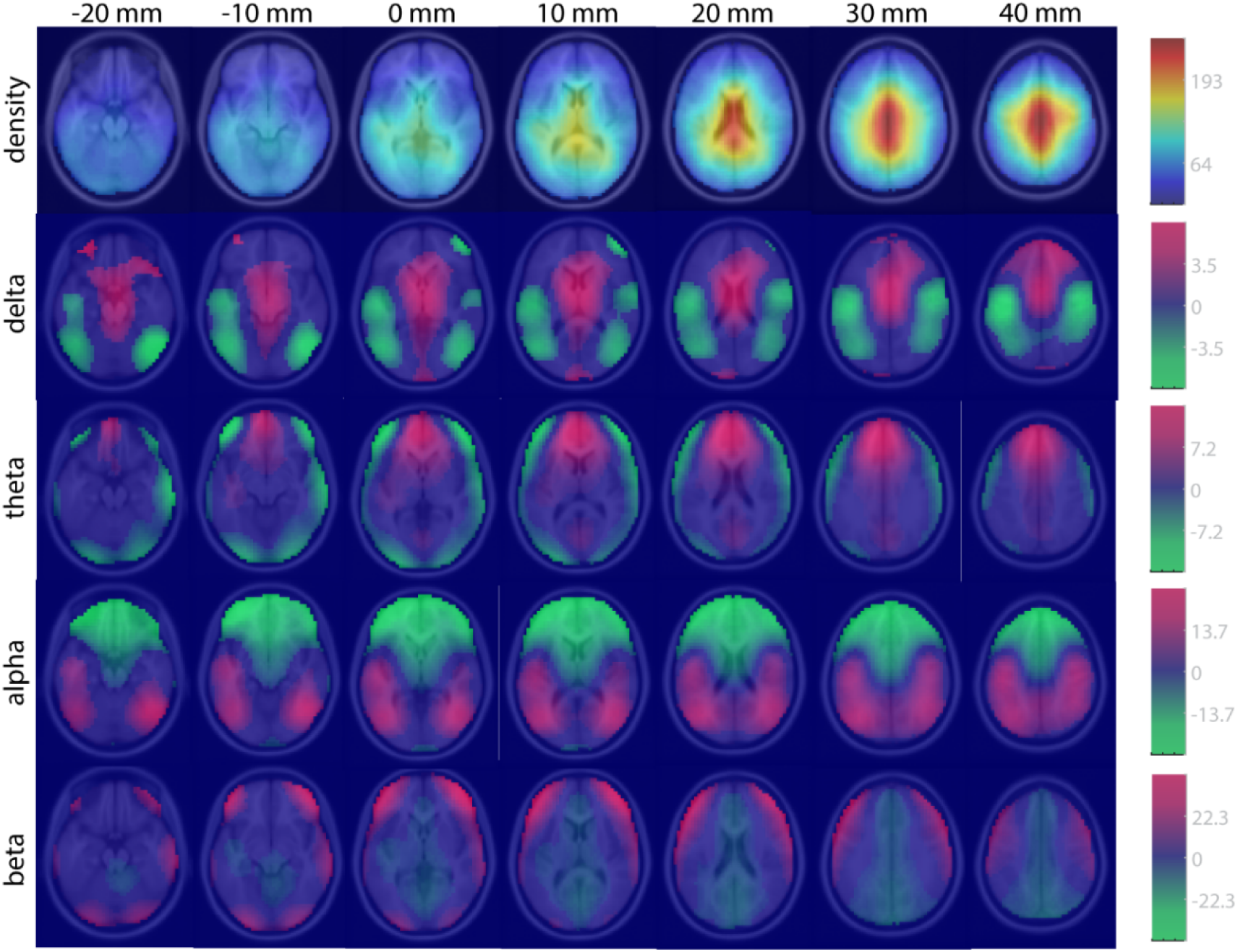
Statistically significant variations of spectral amplitude in common frequency bands. Dipole density plot using 12 mm spatial Gaussian smoothing (top row). Spatially smoothed spectral amplitude deviation percentage for different frequency bands (rows 2-5). Red is associated with higher-than-average baseline amplitude at that frequency and green is associated with lower than average baseline amplitude. The images are masked for significance (*p* < 0.01, *FDR*-corrected).

The bottom four rows of Figure 8 show slice displays of the spatially smoothed amplitude deviation distribution in standard frequency bands. These values are masked by significance (*p* < 0.01, *FDR* corrected), computed by permuting spectral amplitude deviations in each frequency band across dipoles (400 permutations, allowing substitution) followed by spatial smoothing (12 mm Gaussian kernel) to obtain values associated with the null distribution at each brain location. The slice views reflect the distinctive patterns shown in the 3D visualizations of Figure 7.

As in Figure 7, increased beta activity is observed in temporal, lateral frontal, and sensorimotor regions as well as inferior occipital and cerebellar regions, while activity is decreased along the central midline and cingulate structures. Alpha is increased in posterior regions and decreased in anterior regions, while theta is increased in superior frontal and anterior cingulate regions. The fact that regions of increased theta and delta are more prominent in Figure 8 than in Figure 7 indicates that, while significant, the differences for theta and delta are small.

The statistical tests represented in Figure 8 examine whether, for each frequency range, the local amplitude deviation average at a given voxel has a value that is significantly higher or lower than zero. We also performed the same analysis for each study individually and noticed that although the basic patterns were common to most studies, there appeared to be some variations between the studies focusing on visual target detection and those focusing on driving activities.

To examine this variation more carefully, we analyzed two sub-groups that together comprised a large subset of the overall corpus, an *RSVP* group and *Lane-keeping* group. The *RSVP* group (RSVPB, RSVPC, RSVPE, RSVPI, and RSVPU) consisted of studies with different types of visual stimulus presentation within the RSVP paradigm. The recordings in this subset consisted of periods of rapid visual stimulus presentation interspersed with frequent resting periods to alleviate subject fatigue. The *Lane-keeping* group (LKBase and LKCal studies) contained relatively few interspersed events. The right column of Figure 7 shows the spectral amplitude deviations across different frequency bands for these two groups. Although some of the general characteristics of the two groups are similar, there appear to be some differences in detail, particularly for alpha in occipital regions, and for beta in the temporal regions. To test whether these spatial differences were statistically significant, we performed permutation tests to assess whether, at each voxel, the local average spectral amplitude deviation for the *RSVP* group minus that of the *Lane-keeping* group was significantly different from zero. The differences were not significant after controlling for multiple comparisons using *FDR*, indicating a very weak effect, if any.

#### 3.2.2 Some brain areas consistently have higher or lower aperiodic slope

To investigate whether there are systematic variations in aperiodic slope across studies and brain areas, we applied *Fooof* to calculate aperiodic slopes of the “good” dipoles. These are dipoles (56,575 originally) that are inside the brain (26,175) and have *residual variance* < 0.15 (11,637). We further restrict the analysis to dipoles that are identified as non-artifactual (10,170) by the MARA algorithm (Winkler et al., 2011b), and have a linear fit of the aperiodic spectrum with *R*^2^ > 0.95 (9,825). Figure 9 shows the results of this analysis. The average aperiodic slope was 1.1477 +/− 0.0024, a highly significant deviation (*p* < 10^−10^) from pink noise (1/*f* ^α^, α ≈ 1).

**Figure 9.**
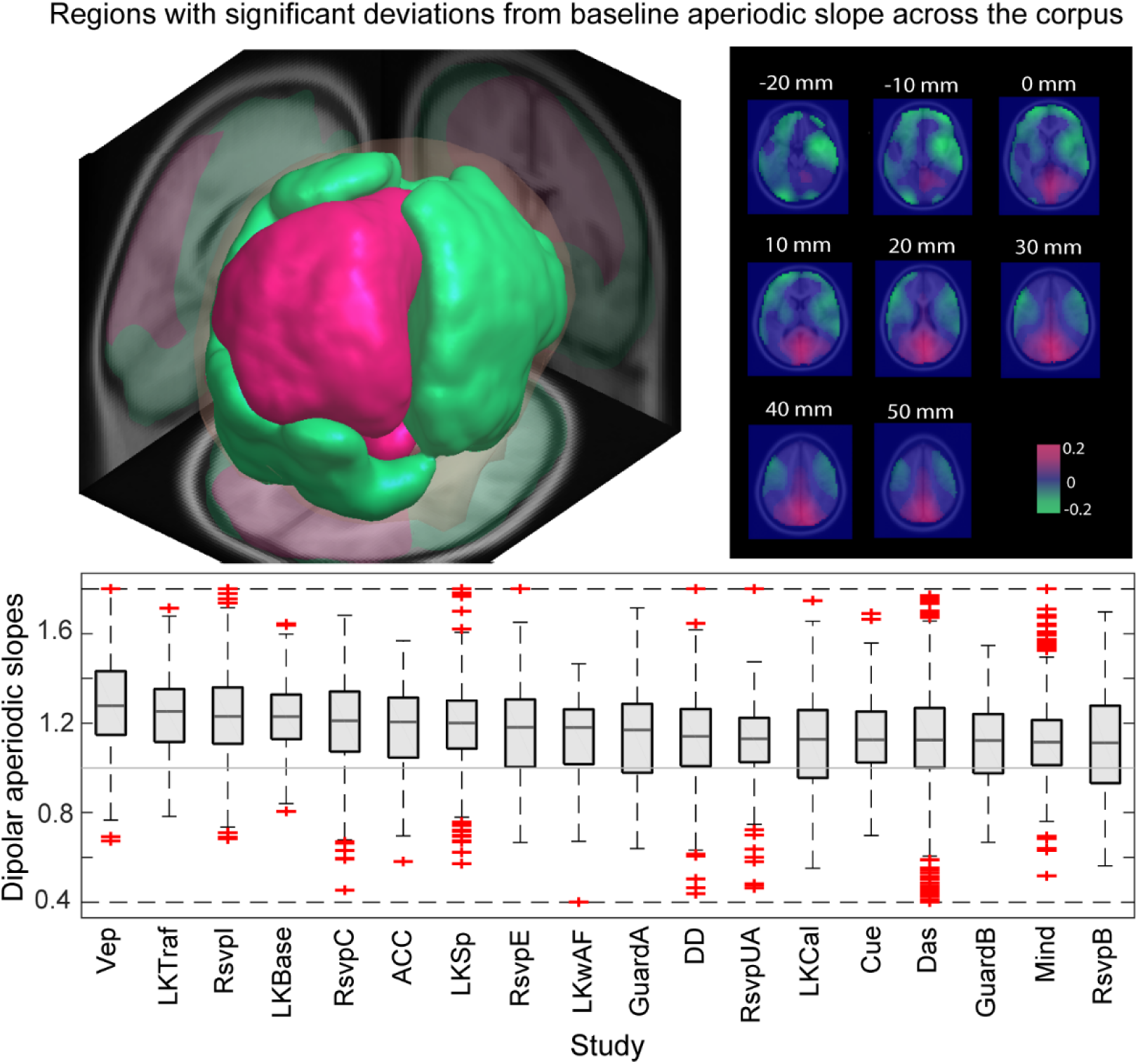
Statistically significant variations of local aperiodic slope. Top left shows three-dimensional view of brain areas with significantly (*p* < 0.05, *FDR*-corrected) high (red) or low (green) local aperiodic slope deviations. Top right shows local average (using 12 mm spatial Gaussian smoothing) aperiodic slope deviations masked by significance. Bottom plot shows distribution of aperiodic slopes by study, sorted in descending order of study median aperiodic slope (based on 9230 dipole power spectra). The gray horizontal line marks a slope of 1.

The top left plot of Figure 9 shows a three-dimensional view of brain areas with significantly (*p* < 0.05, *FDR*-corrected) high (red) or low (green) local aperiodic slope deviations. We obtained this result by applying measure projection analysis (MPA) to the aperiodic slope deviations, mapping the results to brain regions and applying permutation testing to determine statistical significance. The right top plot shows local average aperiodic slope deviations masked by significance. Both left and right temporal areas, plus some occipital areas have lower than average aperiodic slope while posterior midline areas have higher than average slope. The average slope (1.1477 +/− 0.0024) is consistent with the slope for EEG channel data (1.33 +/− 0.19) reported by Dehghani et al. (Dehghani et al., 2010). These authors also report lower slopes in parieto-temporal regions, overlapping with the areas of reduced slope shown in Figure 9.

The bottom graph of Figure 9 shows the distribution of individual dipole aperiodic slopes segregated by study. Studies are ordered in descending order of median aperiodic slope. All of the studies have median aperiodic slope greater than 1 with interquartile ranges roughly centered around their medians. The graph shows that while spectral fall-off varies by dipole, the overall distributions across studies are similar.

## 4 Discussion and conclusions

This paper characterizes EEG channel and source-level signals across a data corpus containing 1,173 recordings representing 633 hours of EEG collected in 18 studies from 6 different institutions on 3 continents using a number of different paradigms and several different headset configurations. Most of the studies fall into two major categories: those with paradigms using visual targets and those with paradigms that include perturbations during driving. While the studies are categorically similar, the experimental details across studies vary considerably, leading to semi-structured information on control variables and subject populations, typical of cross-study comparisons. As public repositories of EEG data become available, questions of how best to compare a new experiment in the context of other published data are an increasingly important. Our results have implications for the aggregation of channel data across studies. In addition, our mega-analysis provides insight into the nominal spatial distribution of single equivalent dipolar sources and their corresponding spectral properties. Limitations of the current corpus include a lack of detailed subject information, such as identification of common subjects across studies conducted at the same institution and a lack of digitized channel locations and/or individual MRI-based anatomical head models to allow for improved covariance analysis and source localization. Nonetheless, the initial analyses and results presented in this paper have implications for the aggregation of channel data across studies. In addition, our mega-analysis provides insight into the nominal spatial distribution of single equivalent dipolar sources and their corresponding spectral properties.

### 4.1 Channel analysis

This study provides some insight into how site/headset/subject/paradigm factors affect EEG signal comparability. One key result of this work is the demonstration that normalizing each recording by a robust recording-specific constant reduces (almost halves) channel amplitude variability across studies. This result is important for any analysis step involving the combination of signals from multiple recordings. Examples include the computation of independent component analysis on datasets formed by combining data from multiple recordings and in various averaging procedures.

Increasingly, BCI (brain-computer interface) applications use data obtained from previous sessions or from other subjects to reduce the amount of calibration data required to train EEG classifiers. Most underlying classifiers assume that training and test signals come from the same distributions or use domain adaption to transform the data to make the distributions similar (Wu et al., 2014) (Wu, 2016). Simple scaling improves the similarity of these distributions. Recently, deep learning using convolutional neural networks has been successfully applied to model EEG signals using raw EEG training data taken from multiple studies (Lawhern et al., 2018), suggesting that as large corpuses of standardized raw EEG data become available, deep learning can potentially lead to advances in EEG analysis similar to those observed in the image and speech recognition domains. Simple preprocessing to improve data distribution similarity across studies may improve results in this area as well.

We have introduced several simple, robust measures for ascertaining whether a given recording is an outlier, whether the study as a whole has the expected behavior, and whether the process for removing eye artifacts has been effective. A scalp map plot of a recording’s amplitude vector provides a simple test of whether a recording is an outlier and whether the eye-artifact removal process has been effective. A simple view of the study as a whole can be achieved by plotting a scalp map of the row medians of the study channel amplitude matrix at different stages of processing (left column of Figure 4). The scalp map of the study channel dispersion vector (Figure 2) plotted at various stages of processing can identify data issues and bad channels across a study. Box plots of study channel dispersion (Figure 3) can also provide useful information about outlier channels and unexpected behavior. Likewise, these methods can be used as part of an evaluation of the effects of different processing methods.

EEG measurements are made through hair, scalp, and skull, which can affect the relative amplitude of channel measurements. However, as shown by the clustering results of Figure 5, relative channel amplitudes may be significantly affected by the EEG cap/sensor technology. These observations suggest that it may be useful to apply channel amplitude normalization techniques before performing group-level or multi-study analyses.

Table 2 shows that normalizing channel data by a recording-specific constant increases the strength of the relationship between channel-derived features (covariance and amplitude) and EEG headset. We can conceptually separate variability caused by EEG headsets into two sources: (1) variability in uniform amplitude/covariance scaling, (2) variability in relative channel features (e.g., channel activity standard deviation ratios). If both of these variability sources were of similar strength, performing a normalization procedure that mitigates type (1) variability should then weaken the relationship between channel correlation/covariance structure and EEG headset type. Since we observe the opposite (an increase in RSA correlation after normalization), we can deduce that the relationship is dominated by variabilities of type (2). Thus, type (1) variability is mostly unrelated to EEG headset type and is reflected as noise within RSA correlations. Future work in this area could explore the potential underlying electromechanical reasons for the phenomenon (e.g., a particular EEG headset design may apply more mechanical force on a subset of EEG sensors, resulting in a better electrical coupling with the scalp). Recently He and Wu (He and Wu, 2018) tested various data-alignment strategies based on covariance matrices. Large-scale testing of data alignment and investigation of signal properties before and after alignment will be an important step in evaluating effectiveness.

Two important caveats are that our results do not explicitly account for the effects of multiple factors on variability, beyond the clear dependence on headset type (displayed in Figure 5). And, secondly, our results are based on full pooling of data (single-level analysis) and do not provide detailed characterization of inferential uncertainty or generalization error within and between recordings, subjects, and studies. Given a suitably structured corpus of data, multi-factor statistical modeling e.g., using ANOVA, can be valuable in characterizing different sources of variability. For example, in a controlled experiment using a multi-factor design, Melnik et al. (Melnik et al., 2017) showed that subject variability dominated differences in event-related potentials, followed by headset type in specific experimental paradigms. Additionally, explicitly modeling recording- and study-specific variability in a multi-level partial-pooling analysis can also help address issues with inflated inferential certainty associated with classical full-pooling analyses. For instance, hierarchical linear modeling (Gelman and Hill, 2006) could be applied within restricted subsets of studies to better characterize effects of various experimental factors on variability. In a companion paper (Bigdely-Shamlo et al., 2018) we apply a more detailed multi-level statistical analysis of a subset of studies from this corpus to characterize the effects of various experimental factors on event-related temporal and spectral features, while explicitly accounting for recording-specific variability.

We use a particular automated pipeline to preprocess the corpus in a standardized way, but the focus of the work is not to establish one preprocessing approach as preferable over another. Rather, the goal of this paper is to show that given a reasonable, consistently-applied preprocessing strategy, EEG mega-analysis is possible and informative. We believe that several automated preprocessing packages such as MARA (Winkler et al., 2011b), AutoReject (Jas et al., 2017), HAPPE (Gabard-Durnam et al., 2018), BEAPP (Levin et al., 2018), and ASR (Kothe and Jung, 2015) (Chang et al., 2018) might be effectively employed in this regard. We have verified that the results of Figure 4 are reproduced using a PREP+MARA pipeline instead of the LARG pipeline.

### 4.2 Spectral analysis

This paper also demonstrated consistent differences in power spectral amplitude within several frequency bands for particular channels and brain regions. Theta increased in superior frontal and anterior cingulate regions. Alpha increased in posterior (occipital, temporal, parietal) regions and decreased in frontal cortex, while delta showed the opposite pattern. Beta increased in temporal, middle frontal and sensorimotor regions and decreased in central midline structures, including superior frontal, superior parietal, posterior cingulate, and inferior occipital regions. We note, however, that we cannot rule out the possibility that the observed beta increases in inferior occipital and temporal regions were influenced by neck or facial muscle activity. Similarly, increased ventromedial prefrontal delta activity could be due, in part, to residual eye movements and blinks.

Modulations in frontal theta and posterior alpha have been well studied and associated with a number of cognitive processes (Ward, 2003). Specifically, decreases in posterior alpha power have been linked to increasing demands in some forms of attention (Klimesch et al., 1998), alertness, and task loading (Holm et al., 2009). Increases in frontal theta power, in surface EEG or sources localized in or near the anterior cingulate and superior frontal gyrus, have been linked to memory encoding (Klimesch, 1999), increased workload (Sammer et al., 2007), cognitive control (Cavanagh and Frank, 2014), and conflict monitoring (Luu et al., 2004).

However, the fact that we did not find statistically significant differences in spectral deviations between the *Rsvp* and *Lane-keeping* corpora suggests that task-related contributions to recording-average IC spectral amplitudes are relatively small compared to other sources of variability such as subject differences. The lane-keeping paradigms are monotonous driving tasks during which prominent resting state network activity has been observed (Lin et al., 2016). Although the RSVP tasks had periods of rapid stimulus presentation, these periods were also frequently interspersed with periods of rest. Also, RSVP presentation rates differed greatly across studies (0.5, 2, 5, 10 or 12 Hz), significantly reducing the frequency-domain effect of visually evoked potentials when spectral averages are computed over all RSVP studies.

Thus, while the patterns of oscillatory activity conform to previously reported spectral phenomena (e.g., frontal theta, occipital alpha), we caution against relating this aggregate spectral activity to specific neural processes or cognitive states such as visual target detection, memory encoding, or fatigue. It is difficult to determine if the regional difference in spectral power are due to the type and number of datasets included in the corpus, systematic limitations in the source analysis, or some universal pattern of brain activity. Rather, these results may be useful as a template or “nominal distribution” from which to measure deviation.

For example, a number of prior studies on lane-keeping tasks have found links between driving performance, fatigue, and spectral power in frontal theta and occipital alpha (Lin et al., 2010) (Borghini et al., 2012) (Chuang et al., 2012). However, findings in this literature have been equivocal as to the direction, magnitude, or consistency of the correlation (Lal and Craig, 2002). Deviations from the nominal distribution may provide a more robust linkage between oscillatory activity, brain state, and task performance. A recent study, using one of the datasets included in this corpus (Garcia et al., 2017) employs a related approach by identifying fluctuations in the relationship between brain activity and behavior (i.e., proactive and reactive brain states) rather than direct correlations between power spectra and performance. These types of analyses, however, require a large, heterogeneous sample in order to establish baseline or nominal states that can generalize across tasks and individuals.

By exploiting a large number of dipoles across studies we reduced the effect of random errors in source localization, but systemic errors, such as the inability of EEG to capture dipoles in certain brain regions, cannot be reduced by simply including more data. We modelled brain sources with ICA-derived single equivalent dipoles and used standard EEG sensor locations and brain templates to localize them. The accuracy of such modelling approaches is a subject of ongoing discussion and investigation (Delorme et al., 2012) (Akalin Acar et al., 2016). We are currently exploring the use of distributed source analysis methods such as LCMV Beamforming (Van Veen et al., 1997), variants of LORETA (Low Resolution Electromagnetic Tomography) (Pascual-Marqui, 2002), and Sparse Bayesian Learning (Wipf and Nagarajan, 2009) (Ojeda et al., 2018) in EEG mega-analysis to address issues with single equivalent dipole modeling assumptions.

The spectral normalization that we are using is similar to relative power spectral density (PSD), but differs in two respects. First, the normalization of a spectrogram that is analogous to a relative PSD involves averaging over time at each frequency, while we take the median in keeping with our robust statistics approach. The second difference is that we are working with spectral amplitude rather than power. Our procedure of dividing the amplitude at each frequency by the Euclidean norm of amplitudes (square root of sum of squared amplitudes over all frequencies) is equivalent to computing the relative power (dividing the squared amplitude by the sum of squared amplitudes), followed by taking the square root of the result. We do not think these differences significantly impact the overall results, but rather make the spectra more comparable across recordings.

The study of aperiodic slope in EEG is relatively new. The aperiodic spectrum has been shown to change with task (Podvalny et al., 2015) and aging (Voytek et al., 2015) (Dave et al., 2018). It is also hypothesized to be associated with tonic differences in excitation/inhibition balance (Voytek and Knight, 2015) (Gao et al., 2017). In a four-subject comparison of EEG and MEG, Dehghani et al. (Dehghani et al., 2010) found spatial variations of the EEG aperiodic spectrum at the channel level that are consistent with the variations found in this study. Brain areas with smaller slope exponent, α, seem to have a large overlap with areas exhibiting significantly higher beta, and vice versa. This relationship suggests the possibility of a common underlying factor inversely relating beta power to aperiodic slope and warrants further investigation.

### 4.3 Conclusion and future work

EEG mega-analysis is a nascent subfield and this work is an initial demonstration of the power and potential for this approach. The automated nature of these analyses and the assembly of a large corpus of data will permit more systematic investigation of brain dynamics associated with cognitive phenomena, along with an assessment of generalizability of results obtained from single studies or paradigms. The metrics introduced in this paper can be used to determine whether particular choices in preprocessing change the general signal characteristics of an EEG dataset or if a particular recording is an outlier for that dataset. The raw data is available upon request through the DataCatalog at https://cancta.net.

In an aforementioned companion paper (Bigdely-Shamlo et al., 2018), we apply mega-analysis with multi-level statistical modeling to assess commonalities in event-related EEG features across studies. Ongoing and future work aims to extend these analyses to conduct more detailed investigations into sources of EEG variability as well as to investigate distributed cortical source activity and large-scale brain connectivity.

## Acknowledgments

The authors would like to thank Scott Makeig for inspiring this work. We would like to acknowledge Tony Johnson and Michael Dunkel of DCS Corporation for their careful assembly and curation of the ARL data. We would also like to acknowledge Ching-Teng Lin and Jung-Tai King of NCTU and the other experimenters who contributed their data to this effort. This work received computational support from UTSA’s HPC cluster Shamu, operated by the Office of Information Technology. This research was sponsored by the Army Research Laboratory and was accomplished under Cooperative Agreement Number W911NF-10-2-0022 (CAST 076910227001). The views and the conclusions contained in this document are those of the authors and should not be interpreted as representing the official policies, either expressed or implied, of the Army Research Laboratory or the U.S Government. The U.S Government is authorized to reproduce and distribute reprints for Government purposes notwithstanding any copyright notation herein.

## Appendix A Data Summary

This appendix provides additional details about the 18 studies used in this paper with references and IRB information. Table A.1 provides some study summary statistics.

**Table A.1:** Additional summary statistics for the 18 studies used in this paper.

| Dataset | Site | Head set | Recs | Subject Info |  | Events | Codes | Mins | Dipoles |  |
| --- | --- | --- | --- | --- | --- | --- | --- | --- | --- | --- |
|  |  |  |  | F/M/U | Ages |  |  |  | Brain | +rv<0.15 |
| Cue | ARLT | B-64 | 33 | 7/10/0 | 19–38 | 297142 | 33 | 1525 | 845 | 403 |
| GuardA | ARLS | B-256 | 29 | 15/12/0 | 20–57 | 211378 | 37 | 1461 | 863 | 221 |
| GuardB | ARLS | B-256 | 22 | 12/9/0 | 19–7 | 169845 | 31 | 1173 | 589 | 120 |
| LKBase | ARLH | B-64 | 21 | 0/5/16 | 27–45 | 1061315 | 35 | 7919 | 3325 | 1068 |
|  | ARLS | B-256 | 81 | 46/34/1 | 19–57 |  |  |  |  |  |
|  | ARLT | B-64 | 28 | 11/17/0 | 18–40 |  |  |  |  |  |
| LKCal | ARLH | B-64 | 21 | 0/5/16 | 27–45 | 518199 | 27 | 3921 | 6581 | 1745 |
|  | ARLS | B-256 | 127 | 55/71/1 | 19–57 |  |  |  |  |  |
|  | ARLT | B-64 | 97 | 48/49/0 | 18–40 |  |  |  |  |  |
| Speed | ARLT | B-64 | 61 | 18/14/0 | 18–40 | 442085 | 26 | 2697 | 1332 | 680 |
| Traffic | ARLT | B-64 | 30 | 12/17/0 | 18–40 | 177937 | 44 | 1389 | 669 | 341 |
| Mind | ARLT | B-64 | 59 | 8/12/0 | 21–38 | 291979 | 50 | 1813 | 896 | 529 |
| RSVPB | ARLS | B-256 | 31 | 16/11/0 | 21–57 | 683108 | 38 | 1970 | 941 | 254 |
| RSVPC | ARLH | B-64 | 51 | 0/0/17 | NA | 82366 | 64 | 392 | 1514 | 292 |
| RSVPE | ARLS | B-256 | 59 | 2/8/0 | 22–51 | 1292141 | 38 | 3652 | 1604 | 462 |
| RSVPI | ARLH | B-64 | 72 | 3/15/0 | 23–58 | 218098 | 45 | 1080 | 2178 | 571 |
| VEP | ARLH | B-64 | 18 | 0/0/18 | NA | 31987 | 14 | 169 | 532 | 160 |
| DAS | NCTU | N-62 | 201 | 46/57/0 | 18–30 | 1219933 | 67 | 5551 | 5152 | 3008 |
| DD | NCTU | N-30 | 60 | 0/15/0 | NA | 203240 | 26 | 1006 | 731 | 536 |
| LKwAF | NCTU | N-30 | 11 | 5/3/2 | 20–30 | 95363 | 14 | 880 | 158 | 145 |
| ACC | TNO | B-64 | 47 | 6/9/0 | 24–60 | 322528 | 57 | 1269 | 1126 | 348 |
| RSVPU | UCSD | B-256 | 14 | 7/1/0 | 19–29 | 484194 | 16 | 980 | 325 | 188 |
| **Total** |  |  | **1173** |  |  | **7802838** | **662** | **38849** | **29116** | **11071** |

### A.1 ARL Auditory Cueing Task (ARL-Cue)

The ARL Auditory Cueing Task involved continuous driving using a driving simulator with brake and foot-pedals but no motion platform. Subjects were instructed to maintain the constant, posted speed and a position in the center of the lane. The vehicle was subjected to perturbations to the left or right at unpredictable times (lane-keeping). In task condition A, a tone was played at random times, unrelated to perturbation events. In task condition B, the tone was played within a 2-second window prior to the perturbation. The two task conditions were counter-balanced among participants and each lasted for 45 minutes. Directly prior to the auditory cueing task, each participant performed 15 minutes of the Calibration Driving Task (A.5). The subject pool consisted of 10 females and 7 males ranging in age from 19 to 38 years old. All participants except one were right handed. This study was conducted by Teledyne Laboratories in Durham, NC using a BioSemi 64-channel headset (Garcia et al., 2017). The study was reviewed and approved by the U.S. Army Human Research Protections Office (Protocol ARL 12-040) before the study began.

### A.2 ARL Advanced Guard Duty Task (ARL-GuardA)

The ARL Advanced Guard Duty Task (Touryan et al., 2016) involved the visual verification of IDs based on eight information fields on an ID card and a corresponding photograph. Participants were asked to determine if the individual in the image, paired with the corresponding ID card, should have access to a specified restricted area. Some of the ID cards were valid and some were not (e.g., expiration date passed, incorrect access area, or photos did not match). Participants were instructed to press either an “allow” or “deny” button for each image-ID pairing. The two-alternative forced-choice response was self-paced with a maximum time limit of 20 seconds. If the participants chose to deny access, they were subsequently asked to provide a reason. Reasons for denied access were selected from a numerical list of five options: incorrect access, expired ID, suspicious DOB, face mismatch, no watermark. If the participant did not respond within the allotted time, the computer forced a “deny” decision. The task was divided into ten blocks of five minutes each. The rate of ID presentation as well as the criteria for validating IDs varied among blocks. The subject pool consisted of 15 females and 12 males ranging in age from 20 to 57 years old. 22 participants were right-handed, 4 participants were left-handed, and one participant was ambidextrous. This study was conducted by SAIC (Louisville, CO) using a BioSemi 256-channel headset. The study was reviewed and approved by the U.S. Army Human Research Protections Office (Protocol ARL 12-041) before the study began.

### A.3 ARL Baseline Guard Duty Task (ARL-GuardB)

The ARL Baseline Guard Duty Task was similar to the ARL Advanced Guard Duty Task. Rate of ID presentation, but not the criteria for verification varied among blocks. The subject pool consisted of 12 females and 9 males ranging in age from 19 to 51 years old. 18 of the participants were right-handed and 3 were left handed. This study was conducted by SAIC (Louisville, CO) using a BioSemi 256-channel headset. The study was reviewed and approved by the U.S. Army Human Research Protections Office (Protocol ARL 12-041) before the study began.

### A.4 ARL Baseline Driving Task (ARL-LKBase)

The ARL Baseline Driving Task was conducted at three different sites using an identical experimental apparatus that included a driving simulator with steering wheel and brake/foot pedals but no motion platform (Touryan et al., 2014) (Brooks et al., 2015) (Brooks and Kerick, 2015) (Garcia et al., 2017). The Baseline Driving Task included either 45 minutes of continuous driving or 60 minutes of driving organized into 10-minute blocks. Subjects were instructed to maintain a center lane position and speed while the vehicle underwent lateral perturbations at random intervals. The subject pool consisted of 43 females, 48 males, and 17 subjects who did not report any demographic information. The subjects ranged in ages from 18 to 57 year old, with 17 subjects not reporting age. 78 subjects were right-handed, 11 subjects were left-handed, 2 subjects were ambidextrous, and 17 subjects did not report handedness. Experiments conducted at ARL HRED (Aberdeen MD) and at Teledyne (Durham, NC) used a BioSemi 64-channel headset. Experiments conducted at SAIC (Louisville, CO) used a BioSemi 256-channel headset. The site protocols were approved by the Army Human Research Protections Office (Protocol ARL-20098-10051, ARL 12-040, and ARL 12-041) before the studies began.

### A.5 ARL Calibration Driving Task (ARL-LKCal)

The ARL Calibration Driving Task consisted of a 15-minute period of continuous driving in which the vehicle was subjected to random lateral perturbations. This task was typically performed at the beginning of experiments that included other, subsequent tasks. The subject pool consisted of 69 females, 59 males, and 17 subjects who did not report demographic information. Subject ages ranged from 18 to 57 years old, with 17 subjects not reporting age. 120 subjects were right-handed, 15 subjects were left-handed, 3 subjects were ambidextrous, and 17 subjects did not report handedness. Like the Baseline Driving Task, these experiments were conducted at three different sites using an identical experimental apparatus that included a driving simulator with steering wheel and brake/foot pedals but no motion platform (Touryan et al., 2014) (Brooks et al., 2015) (Brooks and Kerick, 2015) (Garcia et al., 2017). The ARL HRED site (Aberdeen MD) and the Teledyne site (Durham, NC) used BioSemi 64-channel headsets. The SAIC site (Louisville, CO) used a BioSemi 256-channel headset. The sites protocols were approved by the Army Human Research Protections Office (Protocol ARL-20098-10051, ARL 12-040, and ARL 12-041) before the studies began.

### A.6 ARL Lane-keeping with Speed Control Task (ARL-Speed)

The ARL Lane-keeping with Speed Control Task, which is similar to the ARL Baseline Driving Task, consisted of 45 minutes of continuous driving in a driving simulator while maintaining the posted speed limit. The driving simulator included a steering wheel and brake/foot pedals but no motion platform (Touryan et al., 2014) (Brooks et al., 2015) (Brooks and Kerick, 2015) (Garcia et al., 2017). Subjects were instructed to maintain a constant specified speed and stay in the center of the lane while the vehicle underwent random lateral perturbations. In condition A, speed was controlled by the simulator (i.e., cruise control) while in condition B, speed was controlled by the subject. The Speed Control Task was always counter-balanced with a 45-minute Baseline Driving Task following a 15-minute Calibration Driving Task. The subject pool consisted of 18 females and 14 males, ranging in age from 18 to 40 years old. 27 subjects were right-handed, and 5 subjects were left-handed. The study was conducted at Teledyne Corporation (Durham, NC) using a BioSemi 64-channel headset. The study was reviewed and approved by the U.S. Army Human Research Protections Office (Protocol ARL 12-040) before the study began.

### A.7 ARL Lane-keeping with Traffic Complexity Task (ARL-Traffic)

The ARL Lane-keeping with Traffic Complexity Task, which is similar to the ARL Baseline Driving Task, consisted of 45 minutes of continuous driving in a driving simulator with steering wheel and brake/foot pedals but no motion platform (Touryan et al., 2014) (Brooks et al., 2015) (Brooks and Kerick, 2015) (Garcia et al., 2017). The visual environment consisted of a long, straight highway with visual complexity derived from vehicle traffic, including oncoming traffic and traffic in the direction of travel (in the passing lane). The experiment also included pedestrians on either side of the road, but not crossing the road. Subjects were instructed to maintain a constant specified speed and stay in the center of the lane while the vehicle underwent random lateral perturbations. The 45-minute Traffic Complexity Task was always counter-balanced with a 45-minute Baseline Driving Task following a 15-minute Calibration Driving Task. The subject pool consisted of 12 females and 17 males ranging in age from 18 to 40 years old. 26 subjects were left-handed, and 3 subjects were right-handed. The study was conducted by Teledyne Corporation (Durham, NC) using a BioSemi 64-channel headset. The study was reviewed and approved by the U.S. Army Human Research Protections Office (Protocol ARL 12-040) before the study began.

### A.8 ARL Lane-keeping with Auditory Mind-wandering Distraction Task (ARL-Mind)

The ARL Lane-keeping with Auditory Mind-wandering Distraction Task was similar to the ARL Baseline Driving Task with additional features (Garcia et al., 2017). The experiment was performed using the same driving simulator that included a steering wheel and brake/foot pedals but no motion platform (Touryan et al., 2014) (Brooks et al., 2015) (Brooks and Kerick, 2015). The subject was instructed to maintain lane position and a specified constant speed as the vehicle was subjected to random lateral perturbations. The environment included a mix of regular traffic and police vehicles. The subject was instructed to press a button when spotting a police vehicle. The experiment consisted of three counter-balanced 30-minute blocks corresponding to conditions A, B, and C, respectively. During condition A, the subject listened to an audio podcast on traffic safety; during condition B, the subject listened to a sports-related audio podcast; and in condition C the subject listened to an audio podcast on mediation. The subject pool consisted of 8 females and 12 males ranging in age from 21 to 38 years old. 18 subjects were right-handed, 1 subject was left-handed, and 1 subject was ambidextrous. The study was conducted by Teledyne Corporation (Durham, NC) using a BioSemi 64-channel headset. The study was reviewed and approved by the U.S. Army Human Research Protections Office (Protocol ARL 12-040) before the study began.

### A.9 ARL RSVP Baseline Task (ARL-RSVPB)

The ARL RSVP Baseline Task (Touryan et al., 2014) was a rapid serial visual presentation (RSVP) task utilizing natural photographs containing common objects (chairs, containers, doors, posters, and stairs). A different object was the target for each 10 minute block with a 2-minute break between blocks (6 blocks in total). Images were presented at approximately 5 Hz, and the subject was instructed to press a button each time a target image was perceived. The subject pool consisted of 16 females and 11 males ranging in age from 21 to 57 years old. 22 subjects were right-handed, 3 subjects were left-handed, and 2 subjects were ambidextrous. This study was conducted by SAIC (Louisville, CO) using a BioSemi 256-channel headset. The study was reviewed and approved by the U.S. Army Human Research Protections Office (Protocol ARL 12-041) before the study began.

### A.10 ARL RSVP Cognitive Technology Threat Warning System Task (ARL-RSVPC)

The ARL RSVP Cognitive Technology Threat Warning System Task (Marathe et al., 2014), which was designed to detect neural responses during multitasking, incorporated three simultaneous tasks on different displays (RSVP, target detection, and formation deviation). The center display had a primary RSVP task that presented video clips consisting of five consecutive images, each 100 ms in duration. There was no interval between videos so that the first frame was presented immediately after the last frame of the prior video. Targets included vehicles, animals, and humans, both moving and stationary with varied backgrounds. After receiving a target detection cue, the subject initiates the target detection task on the left display by pressing the “Start” button. The subject sees a set of 20 images selected from the previous RSVP for further evaluation and performs a more detailed assessment by labeling each image as “Target” or “No Target”. After receiving a formation deviation cue, the subject evaluates (vehicle) formation deviation information and indicates the appropriate corrective action to be taken. The cue was presented on the central display (condition 1) or on one of the secondary displays (condition 2). No demographics were reported for the subject pool. The experiment was conducted by ARL HRED (Aberdeen, MD) using a BioSemi 64-channel headset. The study was reviewed and approved by the U.S. Army Human Research Protections Office (Protocol ARL 20098-10025) before the study began.

### A.11 ARL RSVP Expertise Task (ARL-RSVPE)

ARL RSVP Expertise Task (Touryan et al., 2014) used a paradigm similar to the ARL RSVP Baseline Task with images of common objects (chairs, containers, doors, posters, and stairs). A different object was the target for each session. The subject was instructed to press a button each time a target image was perceived. Each subject competed 5 sessions, run on subsequent days. Images were presented at approximately 5 Hz. The subject pool consisted of 2 females and 8 males ranging in age from 22 to 51 years old. 8 subjects were right-handed, 1 subject was left-handed, and 1 subject was ambidextrous. The experiment was conducted by SAIC (Louisville, CO) using a Biosemi 256-channel headset. The study was reviewed and approved by the U.S. Army Human Research Protections Office (Protocol ARL 12-041) before the study began.

### A.12 ARL RSVP Insurgent-Civilian Task (ARL-RSVPI)

The ARL RSVP Insurgent-Civilian Task (Marathe et al., 2015) was an RSVP study with visually similar targets and non-targets presented at a rate of 2 Hz. Target images contained one or more persons holding guns (target), while non-target images contained scenes without people or with people who did not have guns. The experiment was conducted under 4 different RSVP conditions: 1) presentation of only targets with subject counting, 2) presentation of only targets with subject counting and pressing a button for each, 3) presentation of targets and non-targets with subjects counting targets, and 4) presentation of targets and non-targets with subjects both counting and pressing a button for targets. Subjects also performed a baseline condition in which they listed to tones, blinked their eyes in time to audio tones, fixated on a cross, and closed their eyes, each for one minute. The subject pool consisted of 3 females and 15 males ranging in age from 23 to 58 years old. 17 subjects were right-handed, and 1 subject was left-handed. The experiment was conducted by ARL HRED (Aberdeen, MD) using a BioSemi 64-channel headset. The study was reviewed and approved by the U.S. Army Human Research Protections Office (Protocol ARL 20098-09021) before the study began.

### A.13 ARL Visually Evoked Potential Task (ARL-VEP)

The ARL Visually Evoked Potential Task (Hairston et al., 2014) was part of a larger study comparing neural responses for the same subjects on four different commercial headsets and several distinct paradigms. In the VEP task, 18 subjects performed a visual oddball task where target (enemy combatants) and non-target (US soldiers) images were presented at a rate of 0.5 Hz with roughly 1 out of 7 images being targets. Subject information was not available for this study. The experiment conducted by ARL HRED (Aberdeen, MD) using a BioSemi 64-channel headset. This dataset has been used in a number of papers as summarized in a recent Data In Brief (Robbins et al., 2018). The dataset has also been made publicly available on NITRC (https://www.nitrc.org/projects/vep_eeg_raw/). The study was reviewed and approved by the U.S. Army Human Research Protections Office (Protocol ARL 14-042) before the study began.

### A.14 NCTU Dynamic Attention Shifting Task (NCTU-DAS)

The NCTU Dynamic Attention Shifting Task was a lateral perturbation lane-keeping task that was performed on a motion platform and included auditory or visual distractions. On some perturbations, subjects received an attend cue (auditory or visual) along with a direction (left or right). Subjects then received a stimuli (spoken or written word). If the word was a target (animal) subjects pressed the left or right button (corresponding to stimulus position). Subjects heard a “ding” for correctly identified targets and a “buzz” in all other cases. The subject pool consisted of 46 females and 57 males ranging in age from 18 to 30 years old. Handedness information was not available for this study. Data was acquired using a Neuroscan 62-channel headset. The study was approved (VFHIRB no.: 2013-01-029BC) by the Taipei Veterans General Hospital.

### A.15 NCTU Distracted Driving Task (NCTU-DD)

The NCTU Distracted Driving Task (Wang et al., 2015) was a lateral perturbation lane-keeping task performed on a motion platform combined with visual math problems intermittently presented on the dashboard to mimic distracted driving. Subjects pressed a button on the right side of dashboard if the presented equation was correct, and a button on the left side of the dashboard if the presented equation was incorrect. Stimulus onset synchrony was randomly selected to be −400, 0, or 400 ms. Subjects performed four 15-minute blocks separated by 10 minute rest periods. Five conditions included left perturbation (with or without math), right perturbation (with or without math), and math alone. The subject pool consisted of 15 males. Age and handedness was not available for this study. Data was acquired using a Neuroscan 30-channel headset. The study was approved (VFHIRB no.: 2013-01-029BC) by the Taipei Veterans General Hospital.

### A.16 NCTU Lane-keeping with Arousing Feedback Task (NCTU-LKwAF)

The NCTU Lane-keeping with Arousing Feedback Task (Lin et al., 2010) (Huang et al., 2016) was a lateral perturbation lane-keeping task conducted in VR-based driving simulator with perturbations presented at random intervals of between 8 and 12 seconds. Trials that had reaction time 3x longer than trials at the beginning of the session were designated as “drowsy trials”. The alert reaction time was computed as the mean reaction time in the first 5 minutes of the task. In 50% of the “drowsy” trials, the system triggered a 1,750 Hz tone-burst. The subject pool consisted of 5 females and 3 males ranging in age from 20 to 30 years old. These subjects were all right-handed. Two additional subjects provided no demographic information. Data was acquired using a Neuroscan 30-channel headset. The study was approved (VFHIRB no.: 2012-01-019BCY) by the Taipei Veterans General Hospital.

### A.17 TNO Adaptive Cruise Control Task (TNO-ACC)

The TNO Adaptive Cruise Control Task was conducted at the Netherlands Organization for Applied Scientific Research (Brouwer et al., 2017). Subjects drove a Toyota Prius on a closed test track with cruise control set at 35 km/h. An automated human voice indicated whether strong or weak deceleration was desired. The subject then engaged ACC deceleration by pressing a button. Before deceleration a computer voice announced whether the ACC would decelerate strongly or weakly. The ACC decelerated to 25 km/h between 0.5 and 3.5 s after the computer announcement following a steep or a shallow velocity profile (strong or weak deceleration). In 20% of the trials, the deceleration profile did not match the stated desire. The automated human voice then asked the driver to accelerate back to 35km/h. The subject pool consisted of 6 females and 9 males ranging in age from 24 to 60 years old. One subject did not provide age information, and handedness was not available for any of the subjects. Data was acquired using a BioSemi 64-channel headset. The study was reviewed and approved by the U.S. Army Human Research Protections Office (Protocol ARL 16-038) before the study began.

### A.18 UCSD RSVP Target Detection Task (UCSD-RSVPU)

The UCSD RSVP Target Detection Task (Bigdely-Shamlo et al., 2008) used a standard RSVP paradigm with 4.1 second bursts of images presented at 12-Hz. Images consisted of satellite photograph vignettes of London with small airplane targets overlaid. Subjects pressed a button at the end of each 49-image block (left = saw plane, right = didn’t see plane) to indicate whether or not the target was detected. Half of the datasets (train) had “correct/incorrect” visual feedback after button press. The other half of the datasets (test) did not provide feedback. The subject pool consisted of 7 females and 1 male range in age from 19 to 29. Four subjects did not provide age information. Handedness information was not available. Data was acquired using a BioSemi 256-channel headset using a custom cap design. The study was approved (#071254) by the Institutional Review Board (IRB) of the University of California, San Diego.

## Appendix B LARG preprocessing pipeline summary

**Figure B.1.**
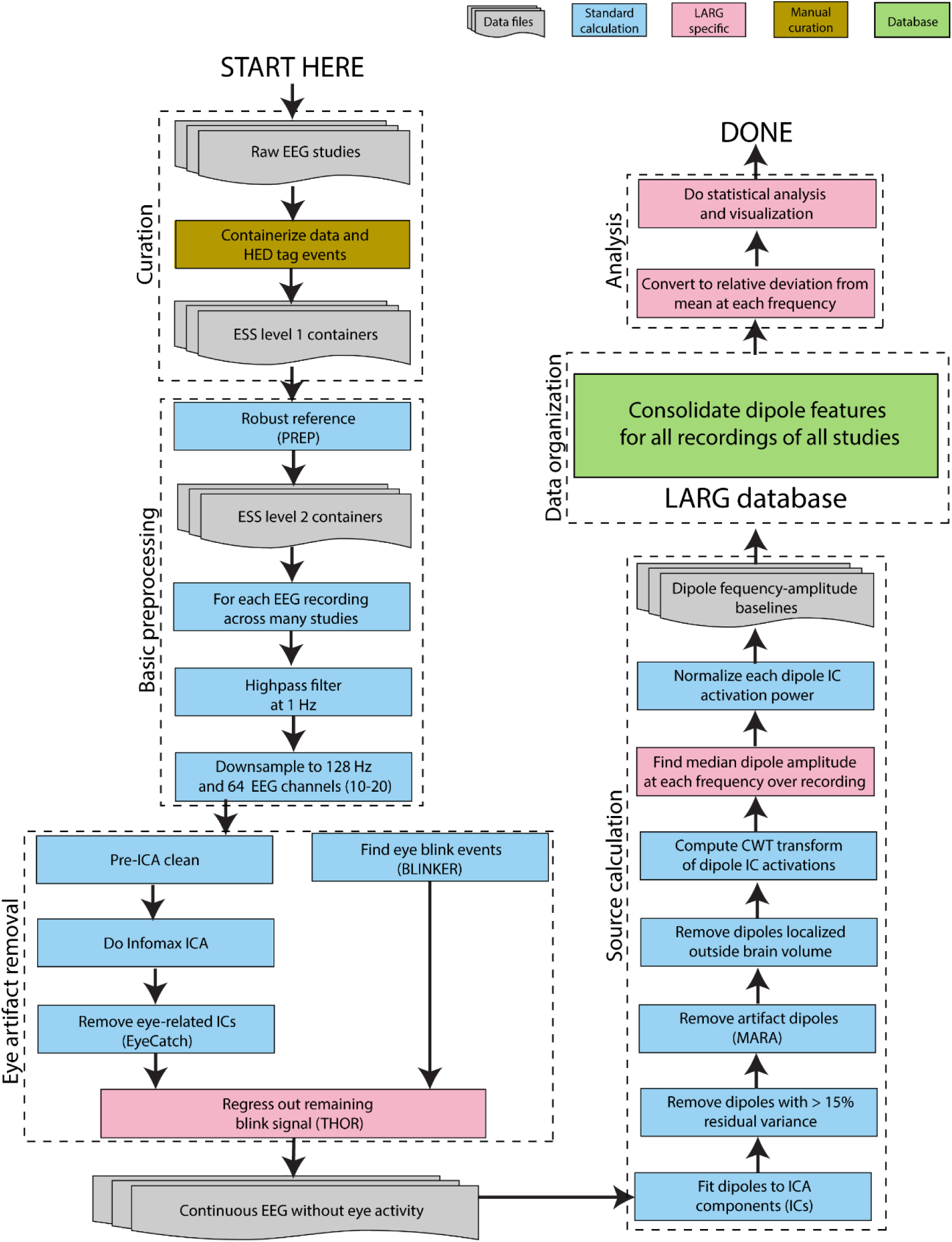
LARG pipeline for automated large-scale analysis of EEG.

## Appendix C Mapping of various headsets to standard 10-20 labels

In combining studies acquired from different headsets, we chose to use the standard positions and labels provided by the manufacturers, rather than individualized headset coordinates (which were not available for most recordings). The reasoning behind this choice was that mapping to a standardized set of coordinates using signal interpolation would introduce significant errors (particularly on the boundaries), which would offset gains in comparability from more exact channel mapping. This appendix illustrates the noticeable differences in channel positions that have resulted from these choices.

Fig. C.1 shows correspondence between channel positions in Biosemi 256-channel (B256) and Biosemi 64 channel (B64) headsets. B256 headsets have detectors that exactly match C3, C4, Cz, Fpz, Fz, Oz, Pz, T7, and T8. The remaining detectors are arranged in concentric circular patterns around Cz. We used a closest Euclidean distance to determine which B256 channel detector was closest to each B64 detector. These associations were unique except in one case where a B64 detector was midway between two B256 detectors.

**Figure C.1.**
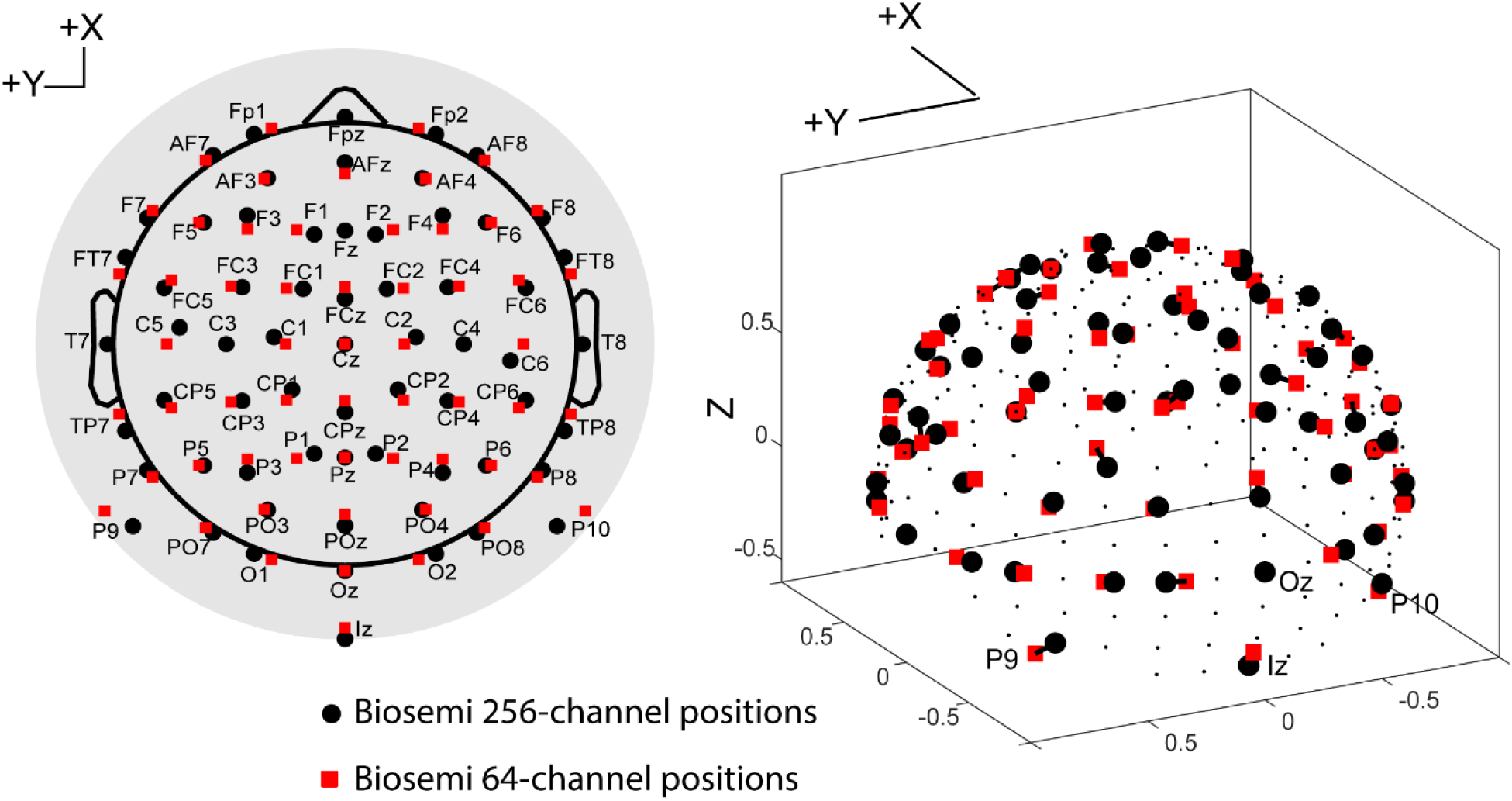
EEGLAB scalp map (left graph) and 3D channel positions (right graph) for two headsets (B256 and B64). Large red squares correspond to B64 channel positions. Large black dots correspond to B256 channels that have been mapped to B64 channels, and small black dots correspond to B256 channels with no corresponding B64 channel.

Fig. C.2 shows the correspondence between Neuroscan 62-channel headset channel positions and Biosemi 64-channel headset channel positions. Although the channel positions appear to agree quite well across the front of the head, there appear to be some systematic differences across the middle and posterior positions. Here detector positions for each headset were scaled so that the average spherical radius of the head was 1 prior to making an association. Closest Euclidean distance then yielded a unique association.

**Figure C.2.**
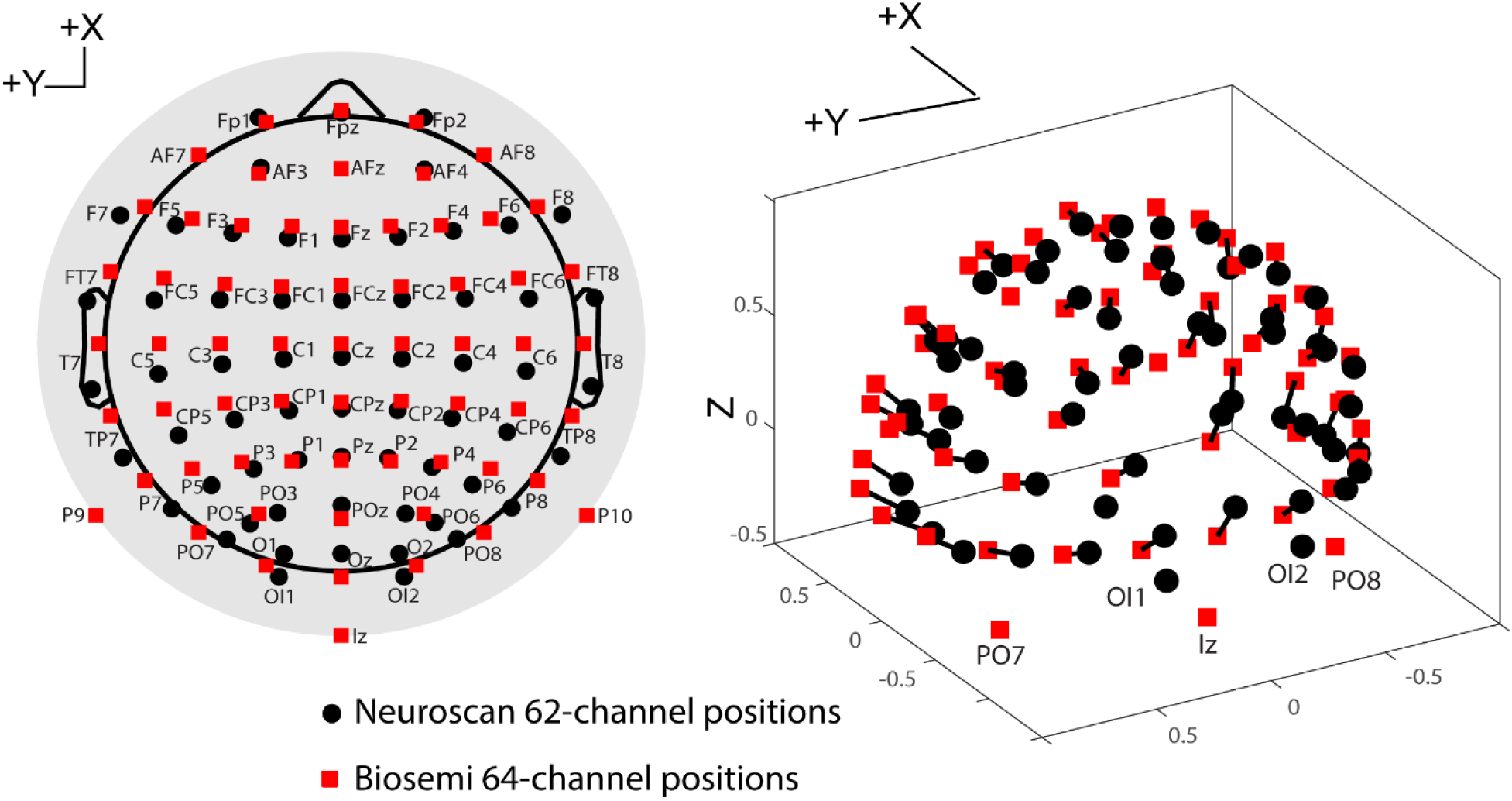
EEGLAB scalp map (left graph) and 3D channel positions (right graph) for two headsets (N62 and B64). Large red squares correspond to B64 channel positions. Large black dots correspond to N62 channel positions.

Fig. C.3 shows correspondence between channel positions in legacy custom 256-channel headset used at the Swartz Center at the University of California San Diego (S256) and Biosemi 64 channel (B64) headsets. The S256 headset had a different channel distribution from the Biosemi headsets, particularly in the lower portions of the head. Closest Euclidean distance did not give good associations for several of the channels in the lower portions of the head. After an initial closest channel mapping was done, the associations were manually adjusted.

**Figure C.3.**
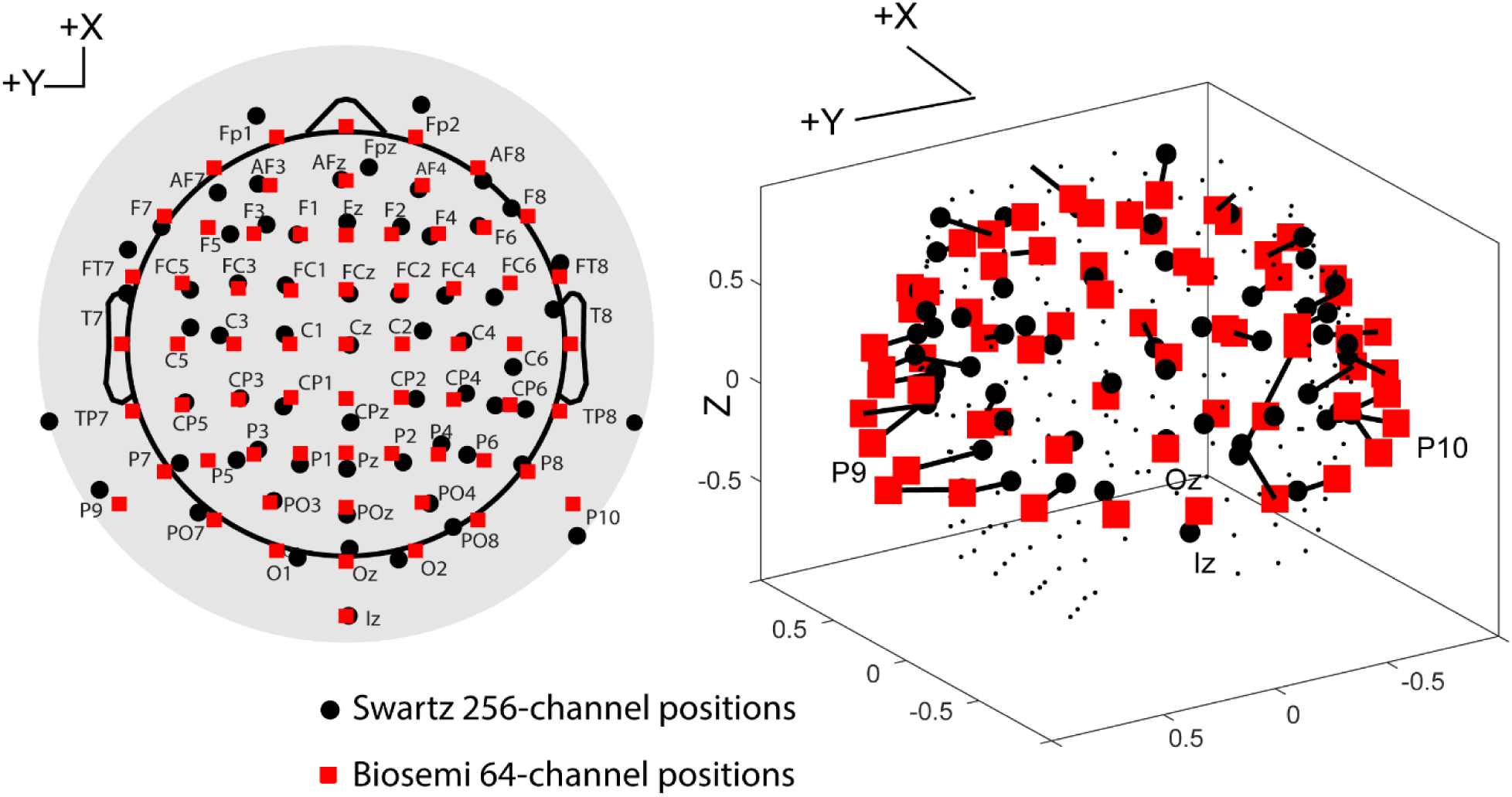
EEGLAB scalp map (left graph) and 3D channel positions (right graph) for two headsets (S256 and B64). Large red squares correspond to B64 channel positions. Large black dots correspond to S256 channels that have been mapped to B64 channels, and small black dots correspond to S256 channels with no corresponding B64 channel.

